# Jasmonate biosynthesis arising from altered cell walls is prompted by turgor-driven mechanical compression and guides root hydrotropism

**DOI:** 10.1101/2020.09.29.319012

**Authors:** Stefan Mielke, Marlene Zimmer, Mukesh Kumar Meena, René Dreos, Hagen Stellmach, Bettina Hause, Cătălin Voiniciuc, Debora Gasperini

**Author notes:** **Corresponding author** Debora Gasperini, Department of Molecular Signal Processing, Leibniz Institute of Plant Biochemistry, Weinberg 3, 06120 Halle (Saale), Germany.

## Abstract

Despite the vital roles of jasmonoyl-isoleucine (JA-Ile) in governing plant growth and environmental acclimation, it remains unclear what intracellular processes lead to its induction. Here, we provide compelling genetic evidence that mechanical and osmotic regulation of turgor pressure represents a key factor in eliciting JA-Ile biosynthesis. After identifying cell wall mutant alleles in *KORRIGAN1* (*KOR1*) with elevated JA-Ile in seedling roots, we found that ectopic JA-Ile resulted from cell non-autonomous signals deriving from enlarged cortex cells compressing inner tissues and stimulating JA-Ile production. Restoring cortex cell size by cell-type-specific KOR1 complementation, by isolating a genetic *kor1* suppressor, and by lowering turgor pressure with hyperosmotic treatments, abolished JA-Ile signalling. Strikingly, heightened JA-Ile levels guided *kor1* roots towards greater water availability, uncovering a previously unrecognized JA-Ile function in root hydrotropism. Collectively, these findings enhance our understanding of JA-Ile biosynthesis initiation, and reveal a novel role of JA-Ile in orchestrating environmental resilience.

## INTRODUCTION

In higher plants, the phytohormone (+)-7-*iso*-jasmonoyl-L-isoleucine (JA-Ile) is a decisive coordinator of growth and stress responses (*1*). Induced JA-Ile signalling secures a successful execution of reproductive development; regulates the acclimation to unfavourable conditions such as drought, salt, cold, elevated ozone and ultra violet light; and is essential to protect plants against insect herbivory, necrotrophic pathogens and RNA viruses (*1-3*). It is estimated that insect herbivory alone impacts >20% of global net plant productivity, and plants unable to synthesize JA-Ile, such as the *allene oxide synthase* (*aos*) mutant of the model plant *Arabidopsis thaliana*, are unprotected against chewing insects (*4, 5*). Rising hormone levels elicit nuclear JA-Ile perception and signalling, leading to the increased expression of JA-Ile marker genes, such as *JASMONATE-ZIM DOMAIN 10* (*JAZ10*), and upregulation of JA-Ile induced defense transcripts such as *VEGETATIVE STORAGE PROTEIN 2* (*VSP2*) and *PLANT DEFENSIN 1*.*2* (*PDF1*.*2*) (*1, 6, 7*). As mounting defense responses are accompanied by reduced vegetative growth, JA-Ile levels are tightly regulated and normally induced only when required (*1, 8*). Despite the broad and critical functions of the jasmonate (JA) pathway in plant growth and environmental acclimation, the intracellular events triggering JA-Ile production remain poorly understood (*1, 9, 10*).

Several molecular elicitors of JA-Ile biosynthesis have been proposed, including herbivore-, microbe-, and damage-associated molecular patterns generated during environmental stresses. These elicitors would be recognized by putative pattern recognition receptors at the cell surface that transduce the signal intracellularly and initiate de novo JA-Ile synthesis, reviewed in (*9, 11*). However, conclusive genetic evidence for ligand-based induction of JA-Ile biosynthesis is still missing. Mechanical wounding is another potent activator of JA-Ile biosynthesis and is widely used to stimulate endogenous hormone production (*6, 10*). In parallel to necrotroph and herbivorous insect attacks, mechanical wounding causes the disruption of cell walls which surround each plant cell and serve as the immediate contact surface with the extracellular environment. Plant cell walls are elaborated structural networks consisting predominantly of complex polysaccharides (cellulose, hemicelluloses and pectins), a lower amount of proteins, and other soluble and phenolic material (*12*). Notably, chemical or genetic inhibition of cellulose biosynthesis by isoxaben application or by mutations in *CELLULOSE SYNTHASE* (*CesA*) genes leads to increased JA-Ile levels, eg. (*11, 13, 14*). Consistently, mutations in *KORRIGAN1* (*KOR1*), a CesA-interacting membrane protein with endo-1,4-beta-D-glucanase activity involved in cellulose biosynthesis, exhibits a slight basal increase in the JA-Ile precursor JA (*15-19*). Nevertheless, it remains unclear how cellulose deficiency stimulates JA-Ile production and in which plant tissues the hormone accumulates.

Plant cells have a high intracellular turgor pressure deriving from a gradient in osmotic potential across the plasma membrane, which is counterbalanced by their cell walls with cellulose microfibrils serving as a load-bearing component (*12*). If osmotic conditions change, plant cells may experience mechanical stress and even deform their shape depending on the extent of the stimulus, their geometry and the properties of their cell walls. Hence, they regulate water and ionic fluxes across the plasma membrane and remodel their cell wall until a new osmotic equilibrium is reached and turgor pressure is re-established (*20, 21*). Cell walls with compromised cellulose microfibrils may be inefficient at counteracting the high intracellular turgor pressure, resulting in enlarged cells. Cell swelling is in fact a typical feature observed in cellulose-deficient mutants or isoxaben-treated plants (*22, 23*). Mechanical stress arising from turgor pressure changes might be therefore involved in activating JA-mediated stress responses (*24*). Interestingly, co-treatments of liquid-grown seedlings with isoxaben and sorbitol serving as osmotica nullified the increased JA levels induced by isoxaben alone (*23, 25*), linking osmoregulation to the JA pathway.

Here, we investigated how cues derived from perturbed cellulose biosynthesis integrate with JA-Ile production. First, we identified *kor1* alleles with increased JA-Ile levels in cell-type-specific contexts of the primary root. While increased JA-Ile signalling in *kor1-4* was confined to inner root tissues, restoring KOR1 function in adjacent cortex cells complemented the JA phenotype. Transversal sections revealed a pronounced enlargement of cortex cells, likely exerting mechanical pressure on physically constrained inner tissues. In fact, reducing cortex cell size by genetic and chemical means fully abolished expanded *kor1* cortex cells and restored JA-Ile signalling, indicating that jasmonate biosynthesis is triggered by turgor-driven mechanical compression. Notably, constitutive JA-Ile signalling did not impact *kor1* growth rate nor its defense responses, but was crucial to guide root growth towards greater water availability.

## RESULTS

### Cellulose-deficient *kor1* mutants have increased root Jasmonate levels

In a forward genetic screen aimed at identifying negative regulators of JA signalling (*6*), we isolated two alleles of *KOR1, kor1-4* and *kor1-5*, exhibiting ectopic expression of the JA-responsive reporter *JAZ10p:GUSPlus* (*JGP*). Basal *JGP* expression in the wild type (WT) as well as in JA-deficient *aos* plants is very weak, whereas *kor1* alleles showed *JGP* activation in the primary root (Fig. 1A, Fig. S1A). Allelism tests with a T-DNA insertion *kor1-6* mutant in the *JGP* background and complementation by transformation with untagged (*KOR1p:KOR1*) and N-terminal CITRINE (CIT)-tagged KOR1 (*KOR1p:CIT-KOR1*) constructs, both restoring the *kor1* short root length, confirmed the causative mutations of the observed *JGP* phenotype (Fig. 1A and B). EMS *kor1* alleles harboured single amino acid exchanges within the extracellular GLYCOSYL HYDROLASE 9 (GH9) domain of KOR1 (L573F in *kor1-4* and P172L in *kor1-5*), while *kor1-6* resulted in lower *KOR1* transcript levels (Fig. S1B and C). Among the three *kor1* alleles, *kor1-5* showed the most severe JA and root elongation phenotypes and *kor1-6* the mildest (Fig.1A and C, Fig. S1A and D), implying that all mutants still retain partial KOR1 function. This is in line with previous reports proposing that full *KOR1* knock-outs are lethal (*16*). Given the milder growth defects and robust JA phenotype of *kor1-4*, we employed this allele for further analyses.

**Fig. 1.**
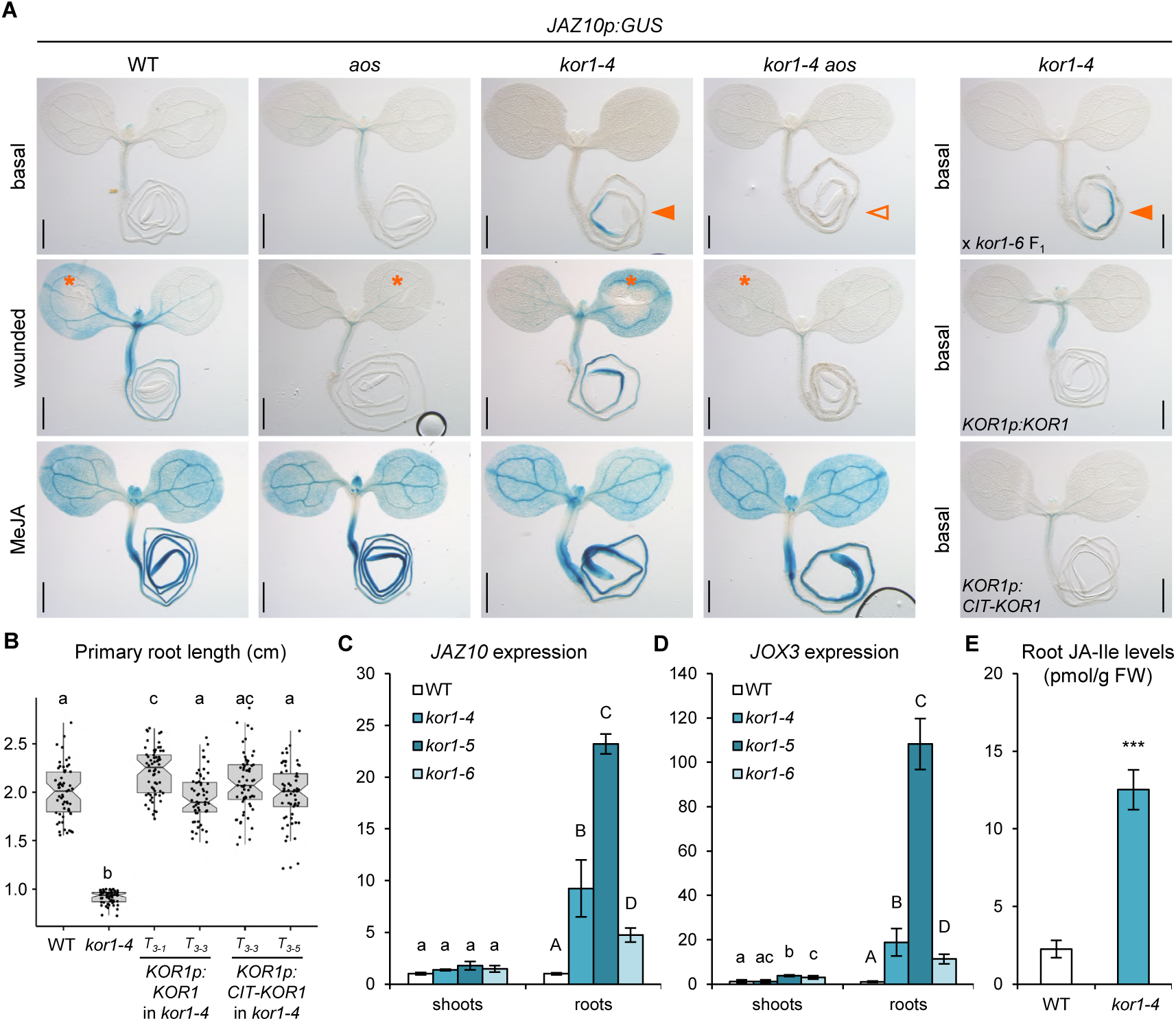
*kor1* mutant roots display increased jasmonate levels and signalling. (**A**) Representative *JAZ10p:GUS* reporter activity in 5-do seedlings of WT, *aos, kor1-4, kor1-4 aos* at basal conditions, 2 h after cotyledon wounding (orange asterisks), 2 h after 25 µM MeJA treatment, and in *kor1-4* x *kor1-6* F_1_ (allelism test), *kor1-4* complemented with *KOR1p:KOR1* and *KOR1p:CIT-KOR1* lines. Note the increased *JAZ10p:GUS* reporter activity in *kor1-4* and allelism test (orange arrowheads), and its absence from *kor1-4 aos* (empty arrowhead). Scale bars, 0.5 mm. (**B**) Primary root length box plot summary in 7-do WT, *kor1-4*, and two independent T_3_ lines for each complementing *KOR1p:KOR1* and *KOR1p:CIT-KOR1* construct. Both constructs restored the *kor1-4* short root phenotype to WT length. Medians are represented inside the boxes by solid lines, circles depict individual measurements (n = 59-61). (**C** and **D**) Quantitative RT-PCR (qRT-PCR) of basal (C) *JAZ10* and (D) *JOX3* expression in shoots and roots of WT and 3 *kor1* mutant alleles. *JAZ10* and *JOX3* transcript levels were normalized to those of *UBC21*. Bars represent the means of 3 biological replicates (±SD), each containing a pool of ∼60 organs from 5-do seedlings. (**E**) Absolute JA-Ile content in WT and *kor1-4* roots. Bars represent the means of three biological replicates (±SD), each containing a pool of ∼600 roots from 5-do seedlings. Letters and asterisks denote statistically significant differences among samples as determined by ANOVA followed by Tukey’s HSD test (P < 0.05) in (B-D), and by Student’s t-test (P = 0.0005) in (E).

Despite the broad *KOR1* expression domain and stunted shoot growth phenotypes of characterized mutants (*26*), constitutive *JGP* activation was not detected in aerial tissues of *kor1* alleles (Fig. 1A, Fig. S1A). Shoot wounding and exogenous MeJA treatment promptly induced *JGP* reporter expression across *kor1* tissues, validating the root specificity of basal *JGP* activity (Fig. 1A, Fig. S1A). Quantitative analysis of *JAZ10* transcripts further supported the specific activation of JA signalling in roots but not shoots of all three *kor1* alleles (Fig. 1C). This phenotype was dependent on JA-Ile production, as increased reporter activity and *JAZ10* transcript levels were abolished in JA-deficient *kor1 aos* double mutants (Fig.1A, Fig. S1A, E and F). In addition to *JAZ10, kor1-4* exhibited increased root expression of JA marker transcripts *JASMONATE OXYGENASE 3* (*JOX3*) and *JAZ3* in a JA-dependent manner, as well as increased levels of the bioactive JA-Ile conjugate (Fig. 1D and E, Fig. S1G). Collectively, our data indicate that KOR1 is a negative regulator of root JA-Ile biosynthesis. Hence, *kor1* mutants represent valuable genetic tools to study how are cell-wall derived signals integrated with intracellular hormone production, and uncover JA-Ile functions in moderating root responses to cellulose-deficiency.

**Fig. S1.**
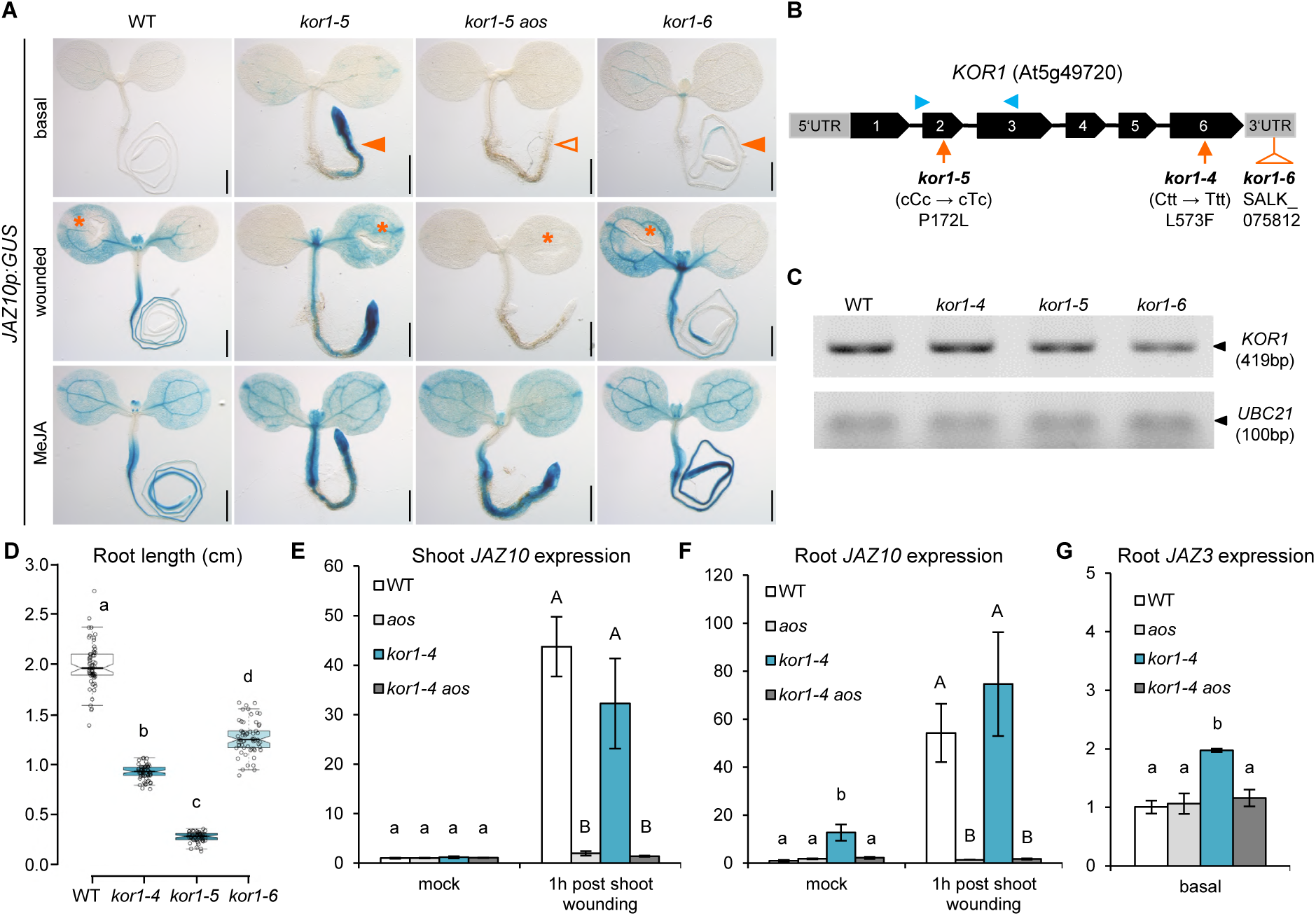
Characterization of JA signalling in *kor1* alleles. (**A**) Representative *JAZ10p:GUS* reporter activity in 5-do seedlings of indicated genotypes basally, 2 h after cotyledon wounding (orange asterisks), and 2 h after 25 µM MeJA treatment. Note the increased *JAZ10p:GUS* reporter activity in roots of *kor1-5* and *kor1-6* mutants (orange arrowheads), and its absence from *kor1-5 aos* roots (empty arrowhead). Scale bars, 0.5 mm. (**B**) Schematic representation of *KOR1* gene structure describing the 3 *kor1* alleles used in this study, and primers (blue arrowheads) used to assess *KOR1* expression by RT-PCR in (C). Black boxes depict exons and lines introns. (**C**) Semi-quantitative RT-PCR of *KOR1* transcripts in WT and 3 *kor1* alleles. Note that *kor1-4* and *kor1-5* are point mutants while the T-DNA insertion in *kor1-6* resulted in a mild reduction in *KOR1* transcript abundance. (**D**) Primary root length box plot summary of 7-do seedlings in indicated genotypes. Medians are represented inside the boxes by solid lines, circles depict individual measurements (n = 51-61). (**E** and **F**) qRT-PCR of *JAZ10* expression basally and 1 h after shoot wounding in (E) aerial organs and (F) roots of WT, *aos, kor1-4* and *kor1-4 aos*. (**G**) qRT-PCR of *JAZ3* basal levels in roots of WT, *aos, kor1-4* and *kor1-4 aos*. Transcript levels in (E) to (G) were normalized to those of *UBC21* and displayed relative to WT controls. Bars in (E) to (G) represent the means of three biological replicates (±SD), each containing a pool of organs from ∼60 5-do seedlings. Letters denote statistically significant differences among samples as determined by ANOVA followed by Tukey’s HSD test (P < 0.05) in (D) to (G).

### Cortex-specific KOR1 expression complements ectopic JA signalling in *kor1* endodermis and pericycle

To map the precise tissues and cell types displaying ectopic JA signalling in *kor1-4* roots, we used a transcriptional reporter expressing three VENUS (3xVEN) fluorescent proteins fused N-terminally to a NUCLEAR LOCALIZATION SIGNAL (NLS) under the control of *JAZ10p* (*JAZ10p:NLS-3xVEN*) (*27*). Similar to *JGP, JAZ10p:NLS-3xVEN* expression was not detectable at basal conditions but increased significantly after MeJA treatment in both WT and *aos* roots, while mechanical wounding triggered *JAZ10p:NLS-3xVEN* induction in the WT but not in *aos* (Fig.2A, Fig. S2). In contrast to the WT, *kor1-4* exhibited constitutive *JAZ10p:NLS-3xVEN* expression predominantly in the early differentiation zone of the primary root, mostly confined to endodermal and pericycle cells (Fig. 2A to C). The extent of activated JA signalling was quantified by evaluating the presence of *JAZ10p:NLS-3xVEN* along longitudinal cell files, and displaying the resulting frequency in a heatmap for each cell layer (Fig. 2D). Weak reporter activation coincided with the onset of cell elongation and proceeded longitudinally into the early differentiation zone for approximately 30 cells before ceasing. While only <10% of *kor1-4* roots showed sporadic *JAZ10p:NLS-3xVEN* expression in a few epidermal or cortex cells, the majority of individuals displayed consistent JA signalling in a stretch of 10-15 endodermal and pericycle cells in the root early differentiation zone (Fig. 2D).

**Fig. 2.**
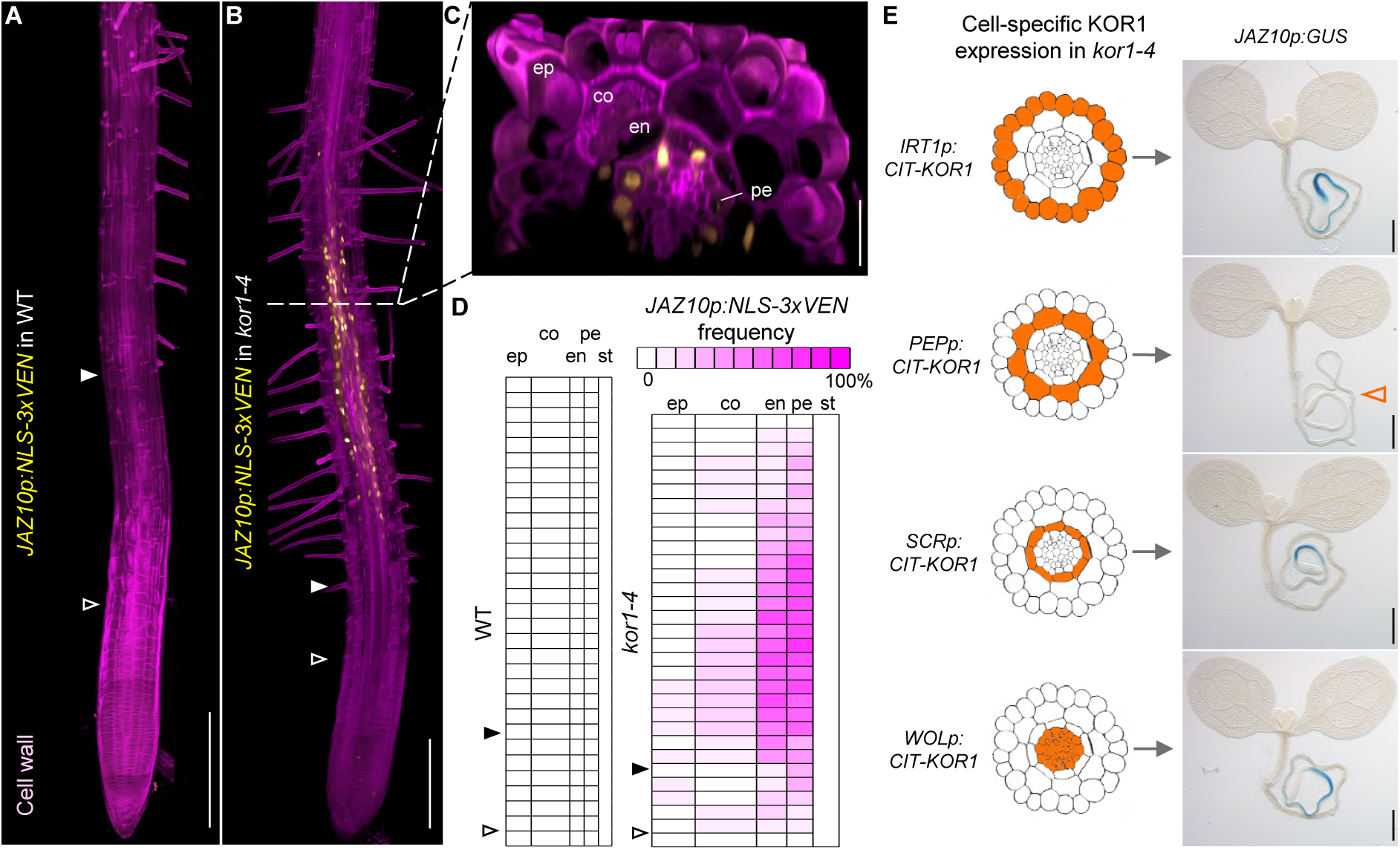
Cortex-specific expression of CIT-KOR1 complements increased JA signalling in endodermis and pericycle of *kor1-4*. (**A** to **C**) *JAZ10p:NLS-3xVEN* expression in 5-do (A) WT and (B and C) *kor1-4* roots cleared with ClearSee, counterstained with the cellulose dye Direct Red 23 and visualized as 3D Z-stacks. (C) Orthogonal view from the epidermis to the vascular cylinder of *kor1-4*. The onset of elongation is indicated by empty arrowheads (first elongated cortex cell), and that of differentiation by filled arrowheads (appearance of root hairs). ep, epidermis; co, cortex; en, endodermis; pe, pericycle. (**D**) Heatmap of *JAZ10p:NLS-3xVEN* frequency in individual cells from WT and *kor1-4* primary roots (n = 21). Presence or absence of the reporter was evaluated from the onset of elongation in individual cells along consecutive longitudinal files for each tissue layer. Reporter expression was not observed in the WT nor in *kor1-4* vascular tissues of the stele (st). (**E**) Cell-specific complementation of *JAZ10p:GUS* activity in *kor1-4* by expressing CIT-KOR1 in either epidermis (*IRT1p:CIT-KOR1*), cortex (*PEPp:CIT-KOR1*), endodermis (*SCRp:CIT-KOR1*) or pericycle and stele (*WOLp:CIT-KOR1*). Note that cortex-expressed CIT-KOR1 complements *JAZ10p:GUS* activity in *kor1-4*, as indicated by lack of the *JAZ10p:GUS* reporter in representative images from T_3_ lines (empty orange arrowhead). Scale bars, 200 µm (A and B), 30 µm (C), and 0.5 mm (E).

Because JA-Ile precursors can relocate across tissues (*28, 29*), increased JA signalling in different cell types could result from cell autonomous or non-autonomous signals. To discriminate between these two possibilities and identify the source tissue responsible for increased JA-Ile biosynthesis, we drove the expression of a functional CIT-KOR1 fusion protein under the control of cell-type-specific promoters and evaluated its capacity to complement the ectopic *JGP* expression in *kor1-4*. As expected, when driven by its endogenous promoter (*KOR1p*), CIT-KOR1 expression was detectable across the entire root, and respective cell-type-specific promoters resulted in CIT-KOR1 expression at the intended locations: *IRON-REGULATED TRANSPORTER 1* (*IRT1p*) in the epidermis, *PLASTID ENDOPEPTIDASE* (*PEPp*) in the cortex, *SCARECROW* (*SCRp*) in the endodermis, and *WOODEN LEG 1* (*WOL1p*) in the stele which includes the pericycle (Fig. S3A to D) (*26, 30*). Intriguingly, the ectopic *JGP* expression in *kor1-4* was complemented only when expressing CIT-KOR1 in the cortex (*PEPp:CIT-KOR1*), but not in the epidermis (*IRT1p:CIT-KOR1*), nor in endodermal (*SCRp:CIT-KOR1*) or pericycle (*WOLp:CIT-KOR1*) cells which exhibited high JA signalling (Fig. 2E). Furthermore, the short root *kor1-4* phenotype was ameliorated by expressing CIT-KOR1 in the cortex, while it remained unchanged when expressing the construct in the epidermis, and it was even slightly exacerbated by cell-type-specific expression in the endodermis or pericycle (Fig. S3E). These results suggest that maintaining KOR1 functionality in the cortex is critical for impeding the activation of JA-Ile production in adjacent endodermal and pericycle cells.

**Fig. S2.**
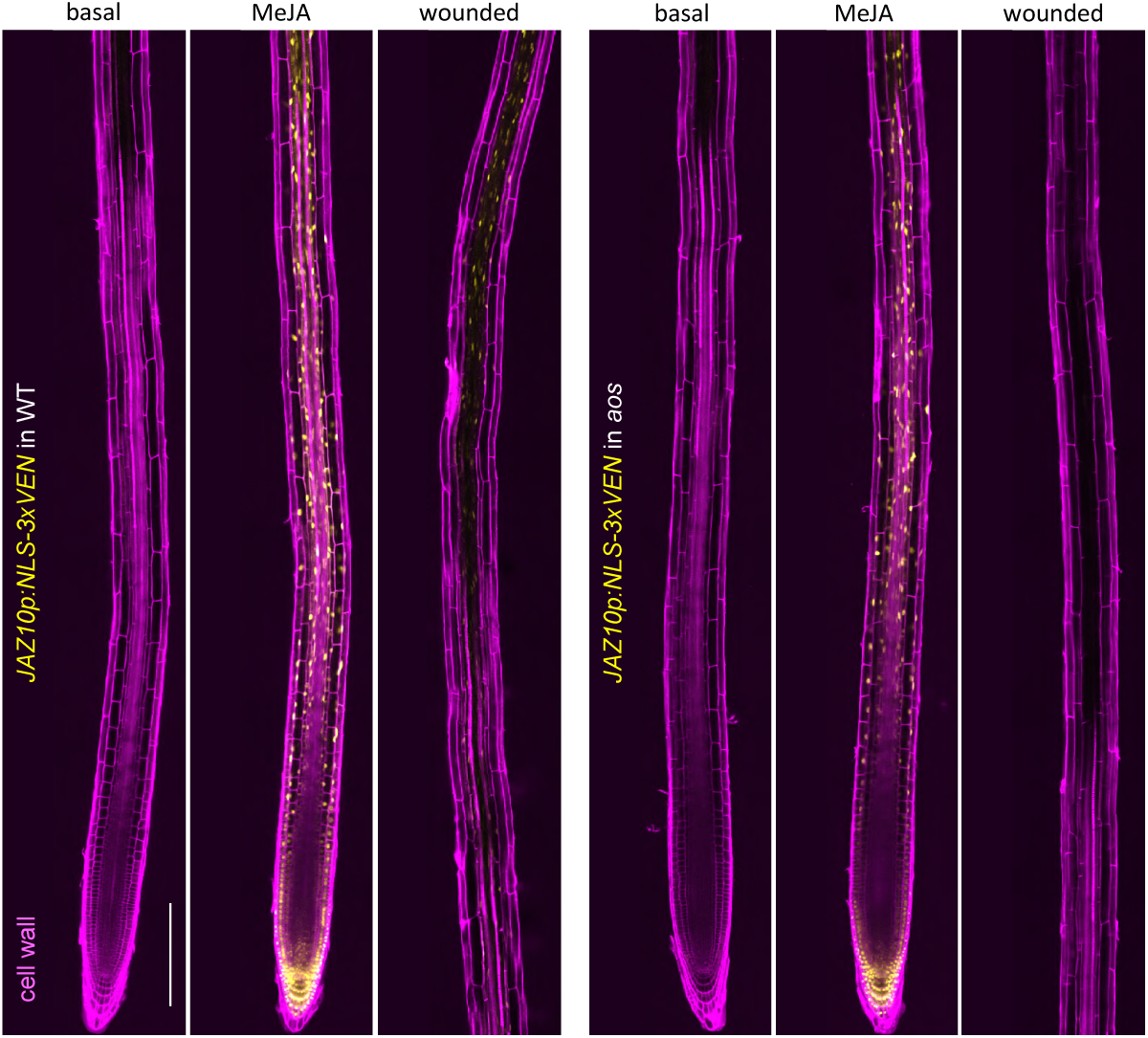
Validation of the *JAZ10p:NLS-3xVEN* reporter. *JAZ10p:NLS-3xVEN* reporter expression in WT and *aos* 5-do roots at mock (basal) conditions, 2 h after 10 µM MeJA treatment and 2 h after cotyledon wounding (n = 10). Cell wall pectins were counterstained with Propidium Iodide. Note that the reporter was activated broadly in WT and *aos* after MeJA treatment while wound-induced activation did not reach the root meristem and was constrained to vascular tissues. Scale bar, 200 µm.

**Fig. S3.**
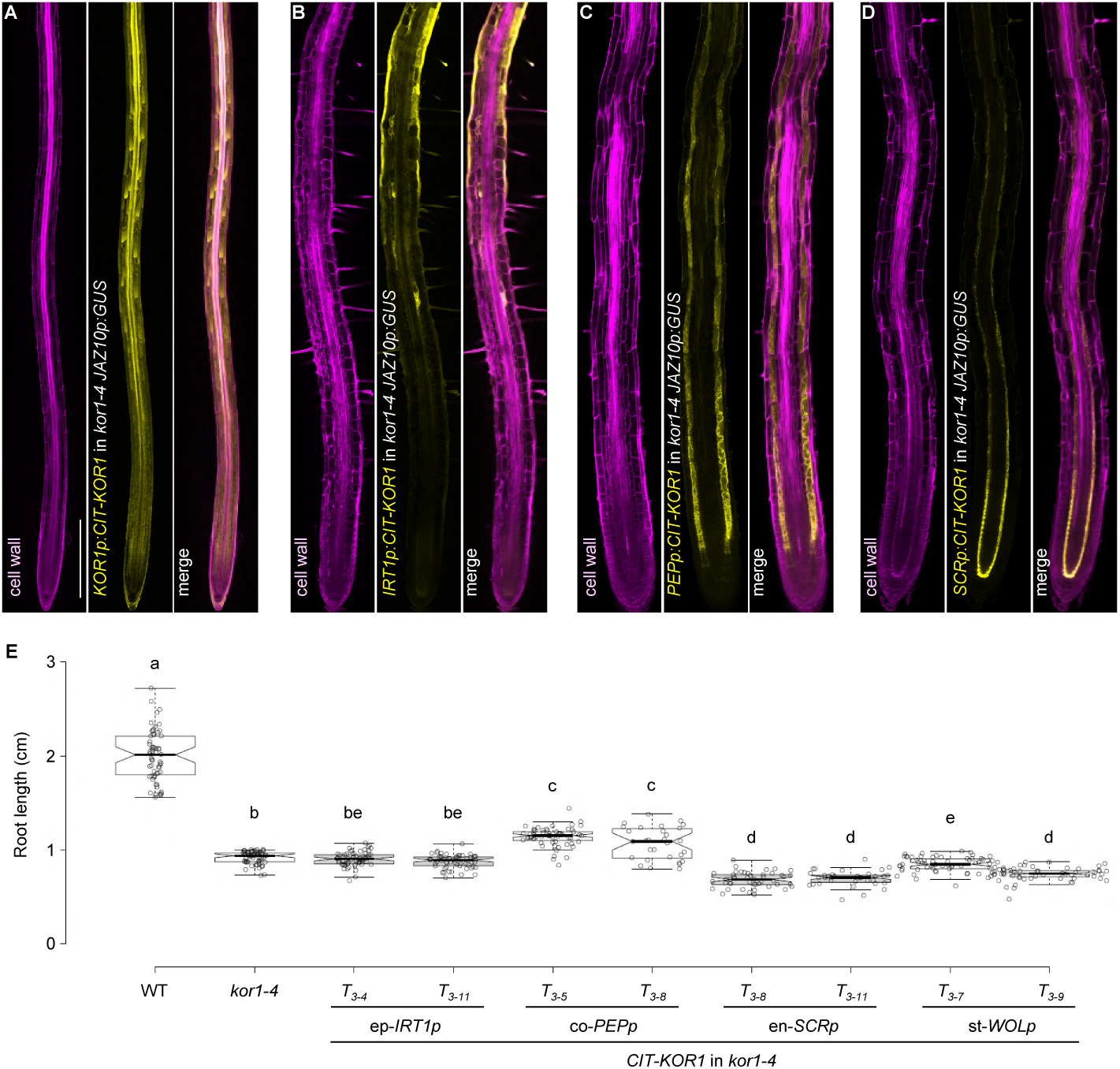
Cell-type specific CIT-KOR1 expression in *kor1-4* and relative root lengths. (**A** to **D**) CIT-KOR1 expression in 5-do *kor1-4 JGP* seedling roots under the control of (A) the native *KOR1* promoter (*KOR1p:CIT-KOR1*), (B) the epidermal *IRT1* promoter (ep-*IRT1p, IRT1p:CIT-KOR1*), (C) the cortex *PEP* promoter (co-*PEPp, PEPp:CIT-KOR1*), (D) the endodermal *SCR* promoter (en-*SCRp, SCRp:CIT-KOR1*). For *WOLp:CIT-KOR1*, we have observed extremely weak CIT fluorescence in pericycle and stele only at higher detector gains. Samples were cleared in ClearSee and counterstained with the cellulose dye Direct Red 23. Scale bar, 200 µm. (**E**) Primary root length box plot summary of 7-do seedlings in indicated genotypes. Medians are represented inside the boxes by solid lines, circles depict individual measurements (n = 30-60), and letters denote statistically significant differences among samples as determined by ANOVA followed by Tukey’s HSD test (P < 0.05). Note that WT and *kor1-4* values are in common with Fig. 1B.

### Mutations in *ESMD1* suppress elevated JA signalling levels in *kor1*

To identify genetic regulators of JA-Ile production in *kor1*, we performed an EMS suppressor screen in *kor1-4 JGP* for lack of JA reporter activation. Mapping-by-whole-genome-sequencing of bulk segregants isolated an allele in *ESMERALDA1* (*ESMD1*), which we named *esmd1-3*. The *esmd1-3* allele abolished the constitutive *JGP* expression and high *JAZ10* levels in *kor1-4* roots while retaining the capacity to induce *JAZ10* transcripts after wounding (Fig. 3A and B, Fig. S4A to C). An ESMD1-mTURQUOISE2(mT) fusion protein expressed under the *ESMD1p* native promoter restored the ectopic JA signalling in *kor1-4 esmd1-3* (Fig. 3A, Fig. S4A). Consistently, introgressing another *esmd1-1* EMS allele (*31*) into *kor1-4* partially suppressed the elevated *JAZ10* levels in *kor1-4* (Fig. 3B), confirming the suppressor’s identity and causative amino acid R373C substitution in *esmd1-3* (Fig. S4D). As we could not retrieve homozygous mutants from a segregating F_2_ *esmd1* population (GABI_216D03), and *esmd1-1* was a weaker JA suppressor than *esmd1-3*, it is likely that full loss of ESMD1 function leads to lethality and that both alleles used herein are partial loss-of-function. *ESMD1* is a member of the plant-specific glycosyltransferase GT106 family, with putative *O*-fucosyltransferase activity involved in regulating pectin homeostasis (*31, 32*). Specifically, *esmd1* was identified as a suppressor of the stunted growth phenotypes of mutants deficient in the major pectin constituent homogalacturonan (HG) (*31*). Because *esmd1* restored the cell-to-cell adhesion defects without complementing the low HG content of pectin mutants *quasimodo1* (*qua1*, a putative galacturonosyltransferase of the glycosyltransferase GT8 family) and *quasimodo2* (*qua2*, a methyltransferase) (*31, 33-35*), we wondered if in addition to suppressing JA signalling, *esmd1-3* was able to revert other *kor1-4* phenotypes.

**Fig. 3.**
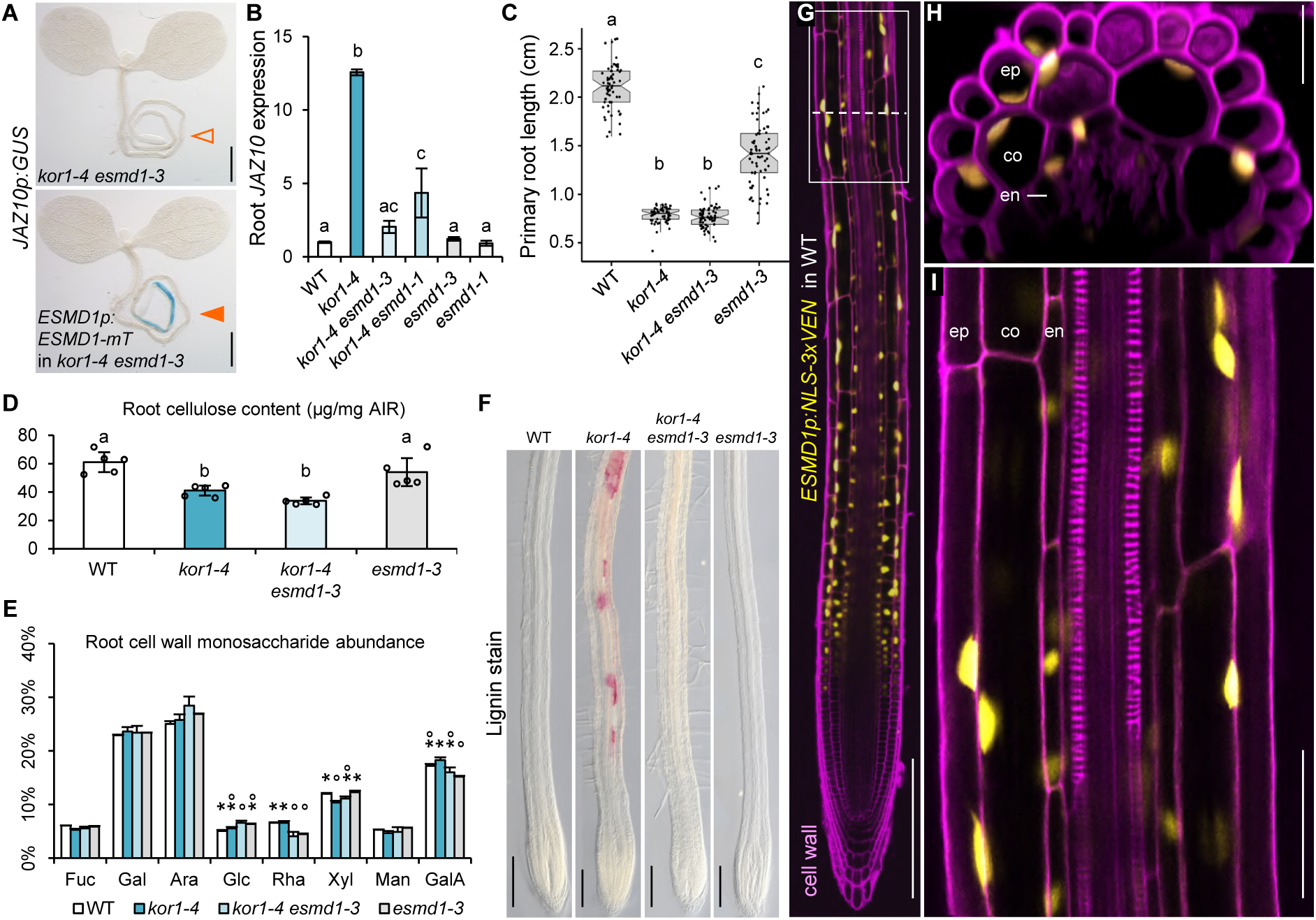
*ESMD1* is a positive regulator of ectopic JA-Ile production in *kor1*. (**A**) *JAZ10p:GUS* expression in *kor1-4 esmd1-3* and its *ESMD1p:ESMD1-mTurquoise*(*mT*) complementation line. Note the lack of *JAZ10p:GUS* reporter activity in roots of *kor1-4 esmd1-3* (empty arrowhead), and its presence in the complemented line (orange arrowhead). (**B**) qRT-PCR of basal *JAZ10* expression in roots of indicated genotypes. *JAZ10* transcript levels were normalized to those of *UBC21*. Bars represent the means of three biological replicates (±SD), each containing a pool of ∼60 organs from 5-do seedlings. (**C**) Primary root length box plot summary in 7-do WT, *kor1-4, kor1-4 esmd1-3* and *esmd1-3* seedlings. Medians are represented inside the boxes by solid lines, circles depict individual measurements (n = 58-66). (**D**) Crystalline cellulose content from alcohol insoluble residue (AIR) extracted from roots of indicated genotypes. Bars represent means of five biological replicates depicted as circles (±SD), each consisting of pools from ∼300 roots from 12-do seedlings. (**E**) Cell wall monosaccharide composition analysis from AIR extracted from roots of indicated genotypes. Bars represent the means of three biological replicates (±SD), each containing a pool of ∼300 roots from 12-do seedlings. Fuc, fucose; Gal, galactose; Ara, arabinose; Glc, glucose; Rha, rhamnose; Xyl, xylose; Man, mannose; GalA, Galacturonic acid. Only statistically significant differences among genotypes indicated with stars and dots assessed for each individual sugar are shown above bars, as determined by ANOVA followed by Tukey’s HSD test (P < 0.001). (**F**) Lignin deposition visualized by phloroglucinol stain (in fuchsia) in primary roots of indicated genotypes. (**G** to **I**) *ESMD1p:NLS-3xVEN* expression pattern in 5-do WT primary roots counterstained with propidium iodide. (H) Orthogonal view from epidermis to vascular cylinder of a Z-stack section in the early differentiation zone through (G, dotted line). (I) Increased magnification in the early differentiation zone from (G, boxed). ep, epidermis; co, cortex; en, endodermis. Letters in (B, C, D) denote statistically significant differences among samples as determined by ANOVA followed by Tukey’s HSD test (P < 0.05). Scale bars, 0.5 mm (A), 200 µm (F and G), 25 µm (H), and 50 µm (I).

Contrary to ameliorating *qua1* and *qua2* growth phenotypes, *esmd1-3* did not restore the short root length nor impacted cellulose content in both shoots and roots of *kor1-4* seedlings (Fig. 3C and D, Fig. S4E). Monosaccharide composition analysis did not reveal major changes in hemicellulose nor pectin constituents between WT and *kor1-4* in both shoots and roots (Fig. 3E, Fig. S4F).

Consequently, and in line with previous reports (*31*), we expected similar neutral and acidic sugar profiles from cell walls of genotypes with *esmd1-3*. Nonetheless, a conspicuous reduction in rhamnose abundance was detected in *esmd1-3* genotypes with respect to the WT, 32-38% in roots and 15% in shoots (Fig. 3E, Fig. S4F). Since cellulose-deficiency often leads to ectopic lignification (*36*), we next evaluated lignin deposition in our mutant set. Phloroglucinol staining revealed patches of lignified cells across the primary root of *kor1-4* that were absent in WT and *aos* plants (Fig. 3F, Fig. S4G). The ectopic lignification in *kor1-4* was not a consequence of elevated JA levels as *kor1-4 aos* still exhibited lignified cells, and was entirely abolished in *kor1-4 esmd1-3*. Overall, the cell wall analysis indicated an elaborated compensatory network in the *kor1 esmd1* double mutant, in which some phenotypes were epistatic to *kor1* (short root length, cellulose deficiency) and others to *esmd1* (lack of increased JA signalling, reduced rhamnose abundance, lack of ectopic lignification). The results therefore suggest that increased JA signalling in *kor1* may be due to indirect consequences of cellulose deficiency, rather than to altered levels of a specific cell wall component.

To gain further insights on how ESMD1 may influence JA responses in *kor1*, we concentrated on its localization in the primary root. Although an ESMD1-GFP fusion protein was expressed in the Golgi when transiently overexpressed in leaf epidermal cells of *Nicotiana benthamiana* (*31*), we were unable to visualize the functional *ESMD1p:ESMD1-mT* construct nor *ESMDp:ESMD1-CIT* in WT Arabidopsis roots, suggesting ESMD1 levels are very low and/or tightly regulated. Instead, *ESMD1p:NLS-3xVEN* expression was mapped to older parts of the root as expected (*31*), and to the epidermis, cortex and endodermis of the early differentiation zone (Fig. 3G to I).

**Fig. S4.**
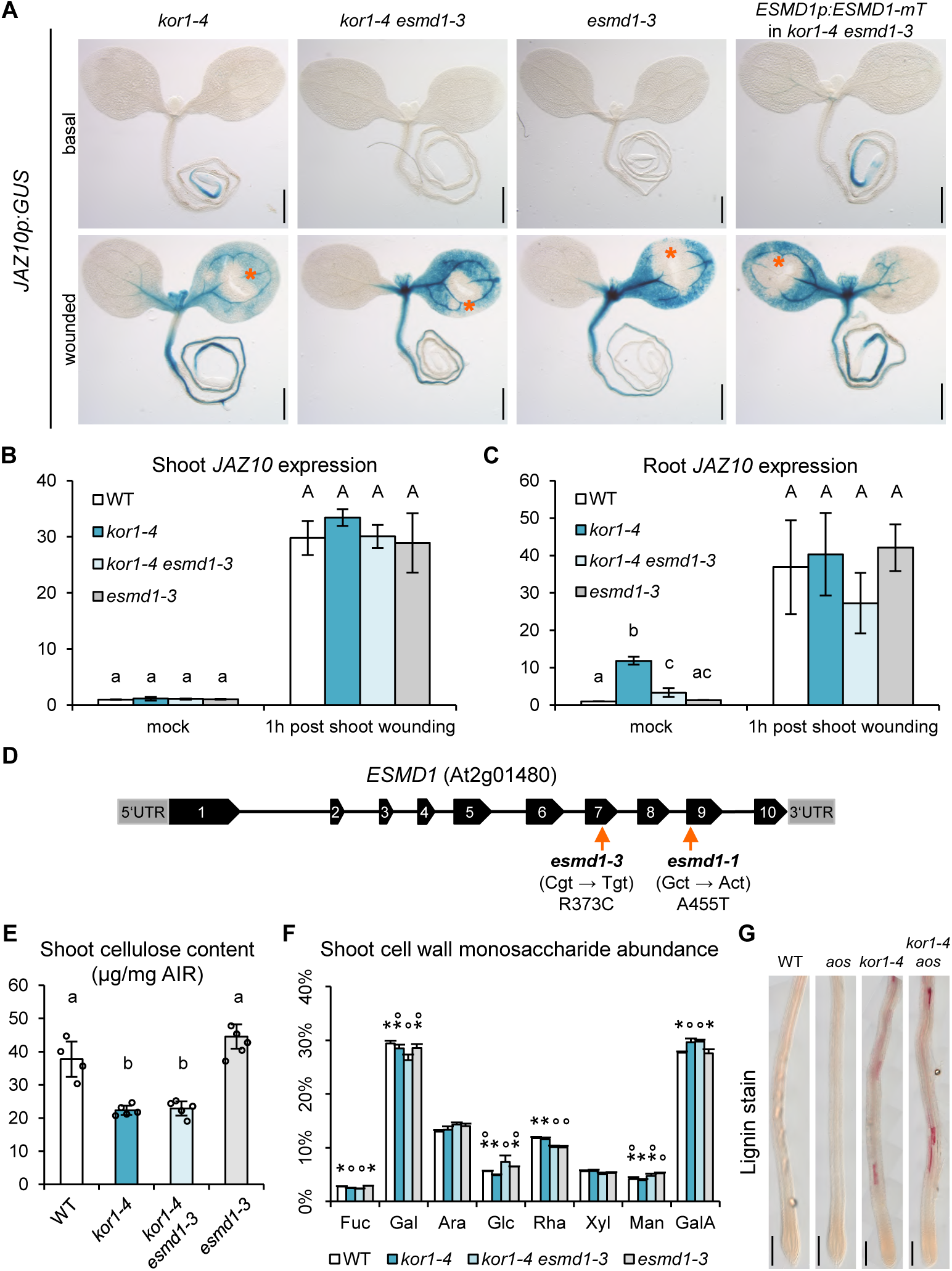
ESMD1 regulates JA levels upon *kor1*-dependent cell wall changes in roots. (**A**) *JAZ10p:GUS* expression in *kor1-4, kor1-4 esmd1-3, esmd1-3* and *kor1-4 esmd1-3* complemented with *ESMD1p:ESMD1-mT* at basal conditions and 2 h after cotyledon wounding (orange asterisks). (**B** and **C**) qRT-PCR of *JAZ10* expression in (B) shoots and (C) roots of indicated genotypes basally and 1 h post cotyledon wounding. *JAZ10* transcript levels were normalized to those of *UBC21*. Bars represent the means of 3 biological replicates (±SD), each containing a pool of ∼60 organs from 5-do seedlings. (**D**) Schematic representation of *ESMD1* gene structure describing the two *esmd1* alleles used in this study. Black boxes depict exons and lines introns. (**E**) Crystalline cellulose content from alcohol insoluble residue (AIR) extracted from shoots of indicated genotypes. Bars represent means of five biological replicates depicted as circles (±SD), each consisting of pools from ∼100 shoots from 12-do seedlings. (**F**) Cell wall monosaccharide composition analysis from AIR extracted from shoots of indicated genotypes. Bars represent the means of four biological replicates (±SD), each containing a pool of ∼100 shoots from 12-do seedlings. Fuc, fucose; Gal, galactose; Ara, arabinose; Glc, glucose; Rha, rhamnose; Xyl, xylose; Man, mannose; GalA, Galacturonic acid. Only statistically significant differences among genotypes indicated with stars and dots assessed for each individual sugar are shown above bars, as determined by ANOVA followed by Tukey’s HSD test (P < 0.001). (**G**) Phloroglucinol stain showing lignin deposition (in fuchsia) in primary roots of indicated genotypes. Letters in (B, C, and E) denote statistically significant differences among samples as determined by ANOVA followed by Tukey’s HSD test (P < 0.05). Scale bars, 0.5mm (A), and 200 µm (G).

### Enlarged *kor1* cortex cells impact JA production in inner tissues

Having found that *ESMD1* expression includes our zone of interest, we further noticed that *kor1-4 esmd1-3* roots are considerably thinner than *kor1-4* (Fig. 4A and B). Transversal sections across the root early differentiation zone coinciding with the sites of JA production in *kor1-4*, revealed that root thickness in the mutant was twice that of the WT and resulted from enlarged areas of all cell-types examined (Fig. 4C and D, Fig. S5A to F). The relatively small expansion of *kor1-4* epidermal cell area (1.1-fold) was counterbalanced by a significant increase in their cell number (Fig. S5A and C). Although aberrant cell divisions were also occasionally observed in *kor1-4* cortex and endodermal cells, the overall cell number in these cell types did not differ significantly from the WT (Fig. S5A). The largest augmentation in *kor1-4* cell area was observed in cortex cells, being 2.6-fold larger than WT, while endodermal and pericycle cells displayed an expansion of 2- and 2.4-fold, respectively (Fig. 4C and D, Fig. S5C to F). These *kor1-4* phenotypes were almost completely restored to WT levels in the *kor1-4 esmd1-3* suppressor, which showed a diminished root thickness, a full complementation of epidermal cell numbers, and reduced cell areas of cortex, endodermis and pericycle cells (Fig. 4A to D, Fig. S5A to F). Notably, the large expansion of *kor1-4* cortex cells was drastically reduced in *kor1-4 esmd1-3* (Fig. 4C and D), leading us to hypothesize that enlarged cortex cells may impinge upon inner tissues.

**Fig. 4.**
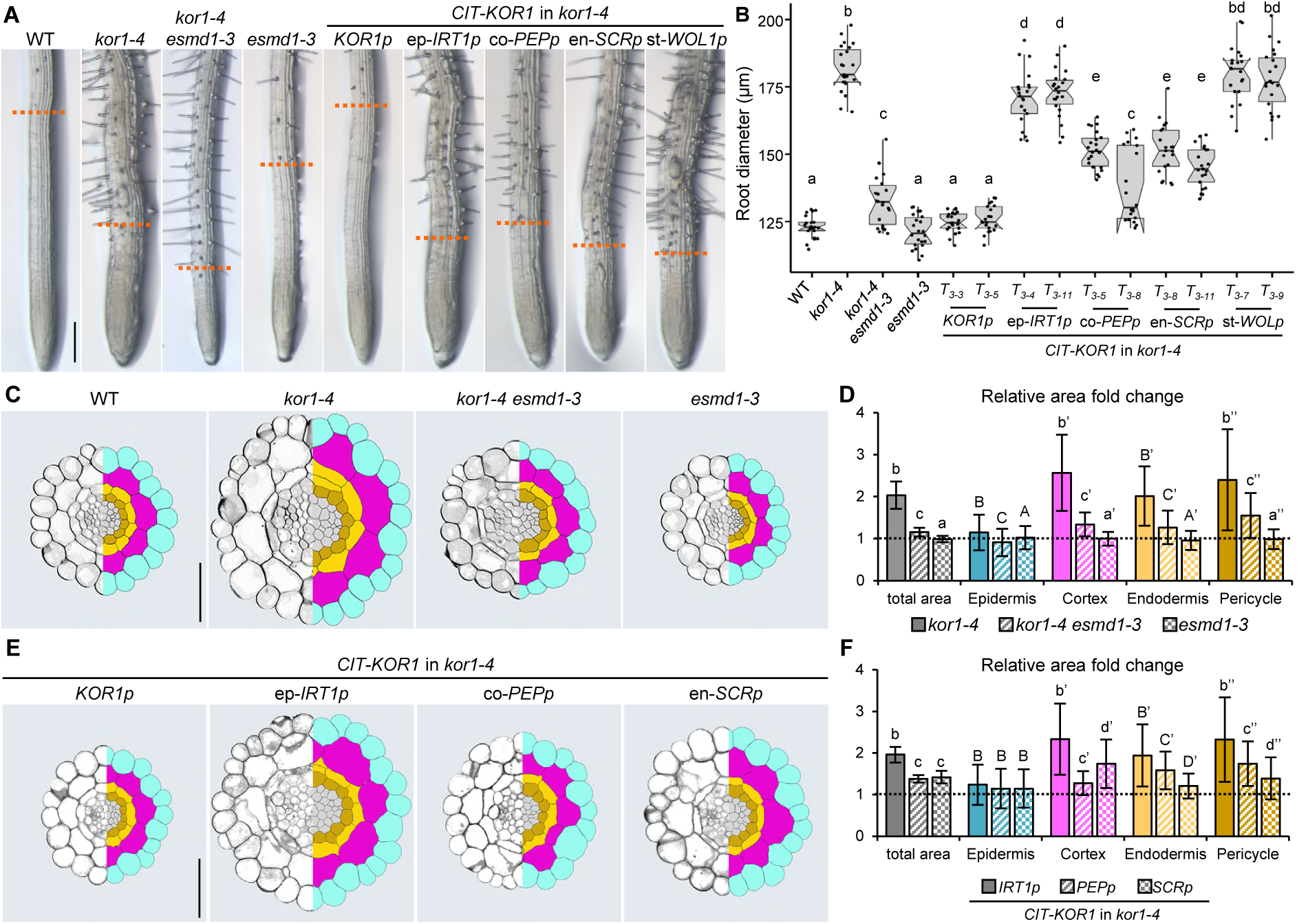
Cortex cell enlargement in *kor1* correlates with constitutive JA production. (**A**) Representative primary root images of WT, *kor1-4, kor1-4 esmd1-3, esmd1-3* and *kor1-4* complemented with CIT-KOR1 under the control of its native *KOR1p* promoter, or epidermis- (ep-*IRT1p*), cortex- (co-*PEPp*), endodermis- (en-*SCRp*) or stele-specific (st-*WOL1p*) promoters. Orange dashed lines define the initiation of the differentiation zone, as indicated by the appearance of root hairs in 7-do seedlings. (**B**) Box plot summary of primary root diameter at the onset of differentiation in 7-do seedlings of indicated genotypes. Medians are represented inside the boxes by solid lines, circles depict individual measurements (n = 22-23). (**C** to **F**) Anatomy and cell size comparisons from transverse sections across the early differentiation zone of the primary root in (C and D) WT, *kor1-4, kor1-4 esmd1-3* and *esmd1-3*, and in (E and F) *kor1-4* complemented with cell-type-specific promoters. (C and E) Representative split images from cross sections (left panels) and respective cell segmentations (right panels). Segmented cell-types are color-coded as: epidermis, turquoise; cortex, magenta; endodermis, yellow; pericycle, mustard; stele, grey. (D and F) Fold change in total and cell-specific areas from segmented transversal root sections in indicated genotypes. Measurements were normalized to those of the (D) WT or (F) *KOR1p:CIT-KOR1*, indicated by dashed lines (individual measurements are available in Fig. S5). Letters in (B, D, F) denote statistically significant differences among samples as determined by ANOVA followed by Tukey’s HSD test (P < 0.05). Scale bars, 200 µm (A), 50 µm (C and E).

To verify if *kor1-4* enlarged cortex cells indeed impacted JA-Ile production in inner epidermal and pericycle, we analysed our transgenic lines expressing CIT-KOR1 under cell-type-specific promoters in transversal root sections coinciding with ectopic JA production (Fig. 2). As expected, expressing CIT-KOR1 under its native promoter fully restored *kor1-4* root thickness to WT dimensions, while epidermis-, cortex- and endodermis-specific CIT-KOR1 expression reduced *kor1-4* root thickness to various degrees. Stele-specific CIT-KOR1 expression (st-*WOLp*) did not have any significant effect and was thus excluded from further analysis (Fig. 4A and B). Compared to the *KOR1p:CIT-KOR1* complemented *kor1-4* line, epidermal CIT-KOR1 expression (ep-*IRT1p*) still exhibited typical *kor1-4* features of 2-fold enlarged root area, increased epidermal cell number, and enlarged areas of cortex, endodermal and pericycle cells (Fig. 4E and F, Fig. S5G to L). Conversely, expressing CIT-KOR1 in either cortex (co-*PEPp*) or endodermal (en-*SCRp*) cell layers, rendered *kor1-4* phenotypes more similar to the *KOR1p:CIT-KOR1* complemented *kor1-4* line by showing a 1.4-fold increase in total root area, and only a milder enlargement of cortex, endodermis, and pericycle cells (Fig. 4E and F, Fig. S5G to L). While the most prominent effect of expressing CIT-KOR1 in cortex or endodermis was to reduce the area of cells in which the fusion protein was localized, the strongest correlation between JA signalling and cell-type area was again found for the cortex cell layer. In fact, expressing CIT-KOR1 in the cortex (co-*PEPp*) abolished *kor1-4 JGP* expression, restored cortex cell area to almost WT levels (1.2-fold) without fully recovering cellular enlargement of endodermal and pericycle cells which persisted being 1.6- and 1.7-fold larger than the *KOR1p:CIT-KOR1* complemented line (Fig. 2E, 4E and F). In turn, endodermal CIT-KOR1 expression (en-*SCRp*) did not abolish elevated JA levels in *kor1-4* nor led to a drastic reduction in cortex expansion which remained 1.7-fold larger, albeit almost completely restoring endodermal and pericycle cell areas (Fig. 2E, 4E and F). As KOR1 activity in cortex cells is important to regulate their size and *JGP* expression in adjacent inner tissues, and *ESMD1* is expressed in both cortex and endodermal cells, it is conceivable that the cortex-endodermis interface is critical for governing JA-Ile production in *kor1*.

**Fig. S5.**
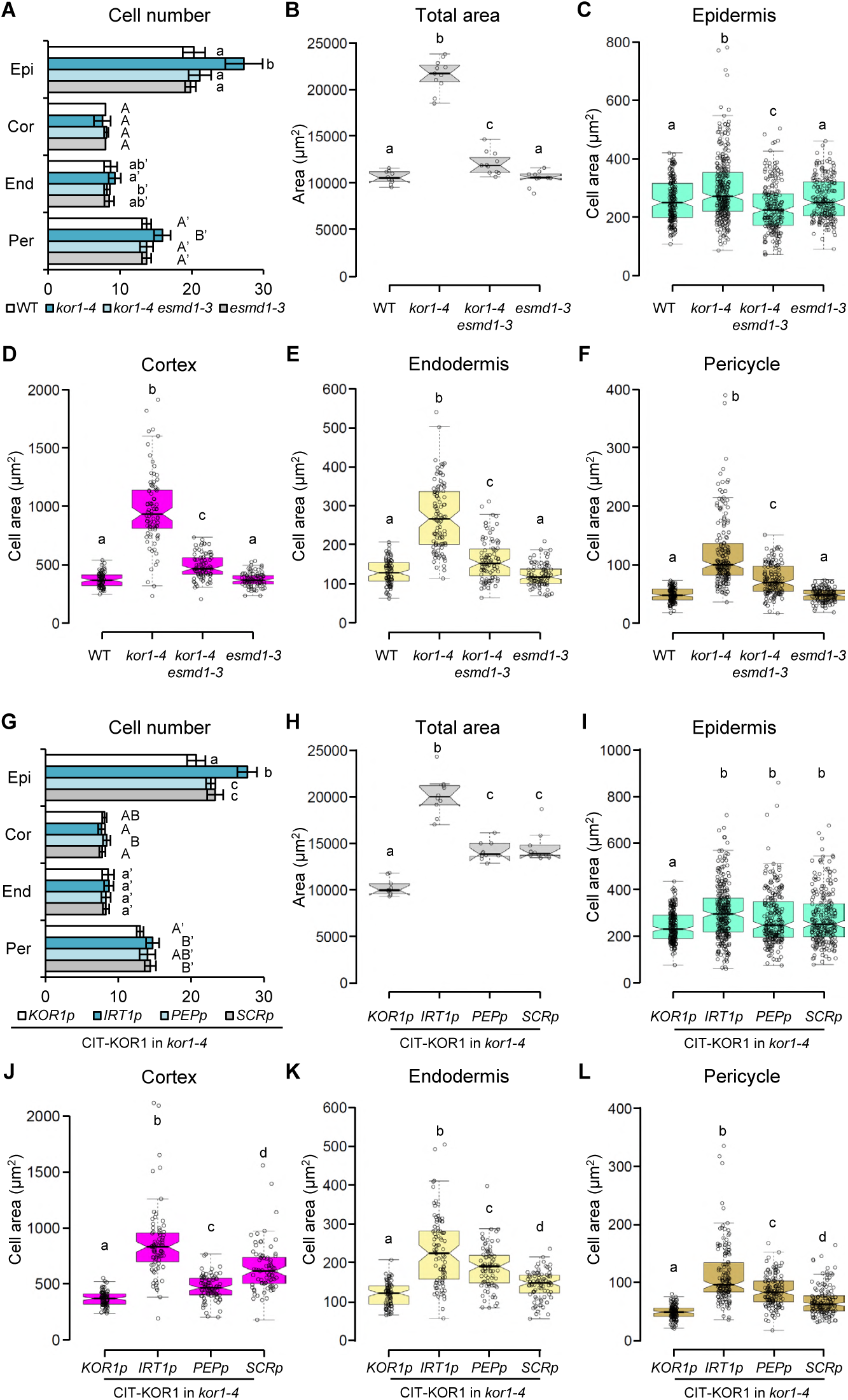
Cellular measurements in transverse root sections of *kor1*, its suppressor *kor1 esmd1*, and its cell-type specific CIT-KOR1 complementation lines. (**A**) Cell number in epidermis (Epi), cortex (Cor), endodermis (End) and pericycle (Per) of the early differentiation zone of primary roots in WT, *kor1-4, kor1-4 esmd1-3* and *esmd1-3* 5-do seedlings. (**B** to **F**) Box plot summary of (B) primary root total area, and cell-type specific areas in (C) epidermis, (D) cortex, (E) endodermis, and (F) pericycle cells from transverse sections of the early differentiation zone in WT, *kor1-4, kor1-4 esmd1-3*and *esmd1-3* 5-do seedlings. (**G**) Cell number in individual tissue layers of the early differentiation zone of primary roots in 5-do *kor1-4* seedlings complemented with *KOR1p:CIT-KOR1* or CIT-KOR1 expressed in epidermis (*IRT1p*), cortex (*PEPp*) or endodermis (*SCRp*) cells. (**H** to **N**) Box plot summary of (H) primary root total area, and cell-type specific areas in (I) epidermis, (J) cortex, (K) endodermis, and (L) pericycle cells from transverse sections of the early differentiation zone in 5-do *kor1-4* seedlings complemented with cell-type specific promoters. Medians are represented inside the boxes by solid lines, circles depict individual cell measurements from 10-11 roots. Letters denote statistically significant differences among samples as determined by ANOVA followed by Tukey’s HSD test (P < 0.05).

### Osmotic support abolishes constitutive JA production in *kor1*

JA responses can be triggered by exogenous treatment with specific pectin- and cellulose-derived fragments serving as ligands binding putative plasma membrane receptors and activating intracellular signalling, reviewed in (*11*). More notoriously, JA-Ile biosynthesis is swiftly activated by mechanical stress (*24*). The lack of a clear association between cell wall composition and JA production in *kor1* (Fig. 3, Fig. S4) led us to hypothesize that increased JA-Ile levels in *kor1* may result from enlarged cortex cells ‘squeezing’ spatially constrained inner tissues, rather than to cell wall-derived elicitors. To test this hypothesis, we decreased intracellular turgor pressure by growing *kor1-4* seedlings under hyperosmotic conditions to withdraw water from their cells. All tested substances acting as osmotica (mannitol, sorbitol, PEG, hard agar) effectively abolished JA signalling in *kor1-4* roots (Fig. 5A and B, Fig. S6A). Mannitol-grown seedlings still responded to wounding, reinforcing the tissue-specificity of the *JGP* reporter (Fig. S6B). Furthermore, while WT root length decreased under hyperosmotic conditions, *kor1-4* root elongation exhibited a tangible amelioration (Fig. 5C). This was also reflected in the reduced area of *kor1-4* transversal root sections grown on mannitol reverting to WT size (Fig. 5D and E). Consistently, mannitol treatment restored *kor1-4* epidermal cell number as well as enlarged epidermal, cortex, endodermis and pericycle cell size (Fig. 5D and E, Fig. S6C to H). Notably, the hyperosmotic treatment fully recovered the enlarged *kor1-4* cortex cell area to WT levels, while endodermal and pericycle cell were still 20% larger with respect to the WT (Fig. 5D and E). Thus, the activation of JA production in *kor1* is consistent with inner tissues being mechanically stressed by enlarged cortex cells.

**Fig. 5.**
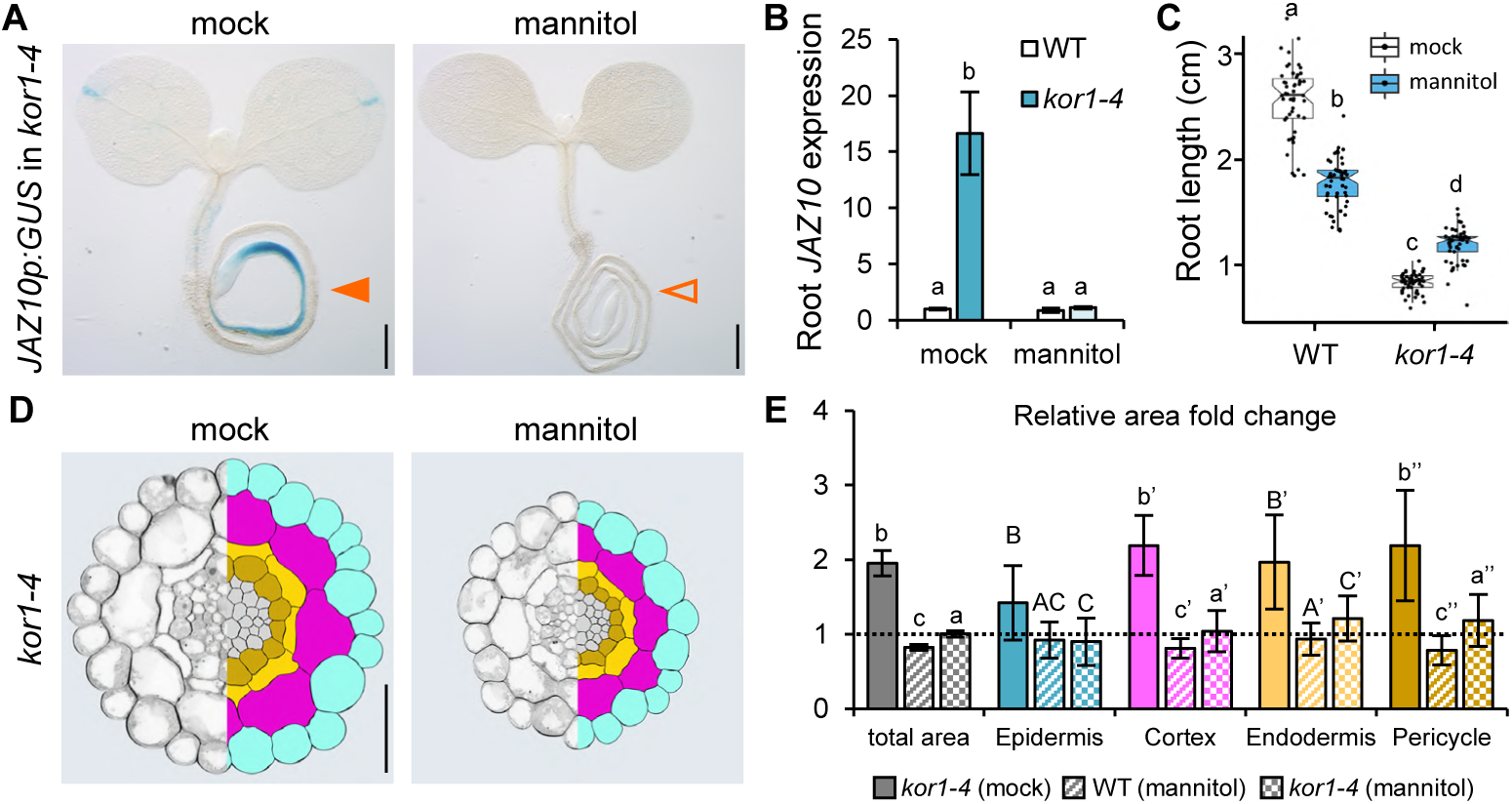
Hyperosmotic mannitol treatment relieves JA signalling in *kor1* to WT levels. (**A**) *JAZ10p:GUS* reporter activity in 5-do *kor1-4* seedlings grown in mock or 3% mannitol-supplemented media. Note the presence of ectopic JA signalling in mock conditions (orange arrowhead) and its abolishment in mannitol (empty arrowhead). (**B**) qRT-PCR of basal *JAZ10* root expression of WT and *kor1-4* grown in the presence or absence of 3% mannitol. *JAZ10* transcript levels were normalized to those of *UBC21*. Bars represent the means of 3 biological replicates (±SD), each containing a pool of ∼60 organs from 5-do seedlings. (**C**) Primary root length box plot summary of 7-do WT and *kor1-4* seedlings grown in the presence or absence of 3% mannitol. Medians are represented inside the boxes by solid lines, circles depict individual measurements (n = 59-61). A linear model was used to assess differences in responsiveness between the two genotypes: WT plants responded with a decrease in root length (−0.78 cm; p-value: 2e-16), whereas *kor1-4* with an increased (+0.35 cm; p-value: 2e-16). (**D**) Representative split images from transverse sections (left panels) and respective cell segmentations (right panels) across the early differentiation zone of primary WT and *kor1-4* roots grown in the absence (mock) or presence of 3% mannitol. Segmented cell-types are color-coded as: epidermis, turquoise; cortex, magenta; endodermis, yellow; pericycle, mustard; stele, grey. (**E**) Fold change in total and cell-type specific areas from segmented transverse sections of WT and *kor1-4* grown with and without 3% mannitol. Measurements were normalized to those of the mock-treated WT indicated by a dashed line (individual measurements are available in Fig. S6). Letters in (B, C, and E) denote statistically significant differences among samples as determined by ANOVA followed by Tukey’s HSD test (P < 0.05). Scale bars, 0.5 mm (A), and 50 µm (D).

**Fig. S6.**
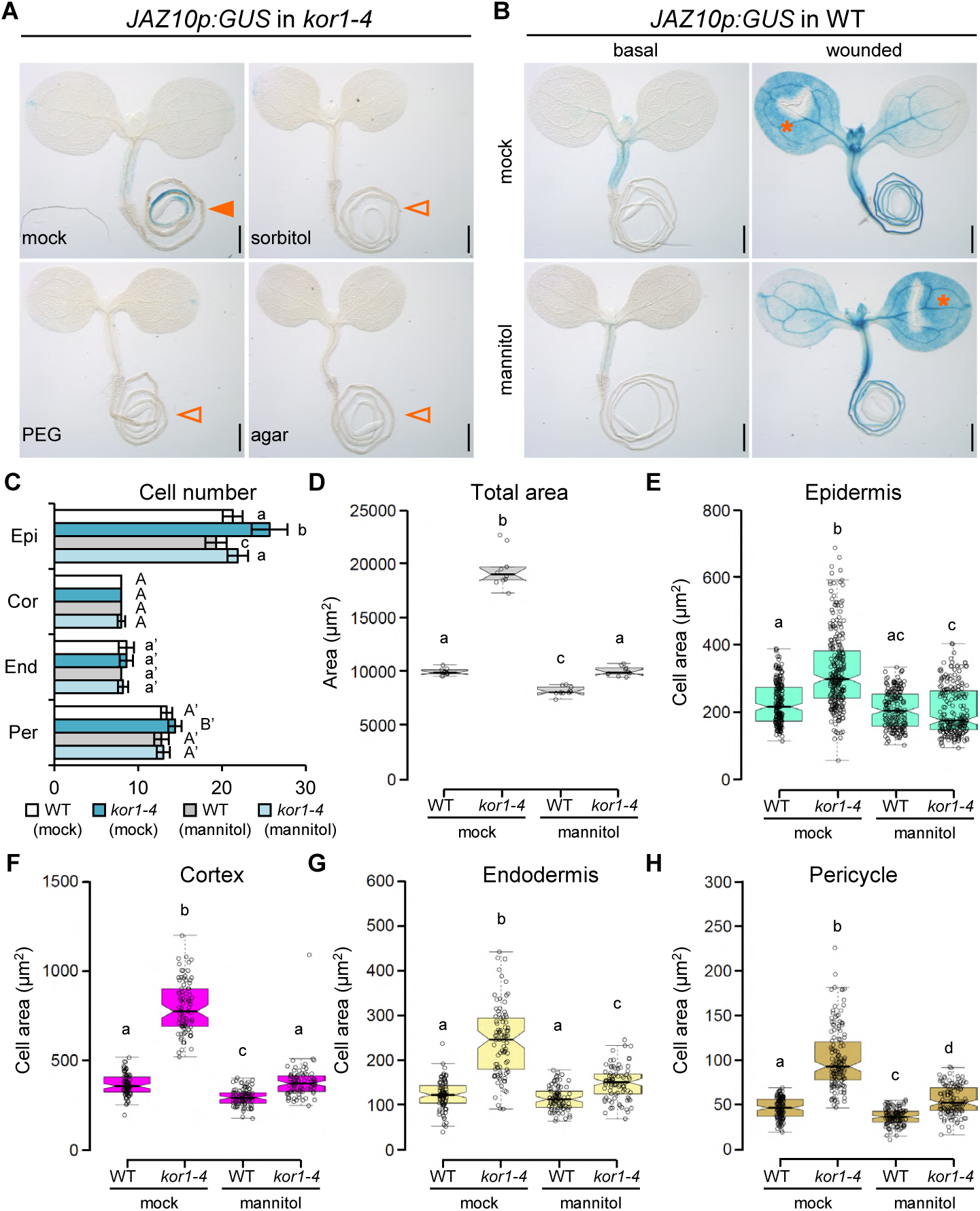
Hyperosmotic treatments and cellular measurements. (**A**) Basal *JAZ10p:GUS* reporter activity in 5-do *kor1-4* seedlings grown in the absence (mock) or presence of 3% sorbitol, 3% PEG or in 3% agar. Note the presence of ectopic JA signalling in mock conditions (orange arrowhead) and its abolishment in hyperosmotic media (empty arrowhead). (**B**) *JAZ10p:GUS* reporter activity in 5-do WT seedlings grown in the absence (mock) or presence of 3% mannitol under basal conditions and 2 h after cotyledon wounding (orange asterisks). Scale bars, 0.5 mm (A and B). (**C**) Cell number in individual tissue layers of the early differentiation zone of 5-do primary roots of WT and *kor1-4* seedlings, grown in the absence or presence of 3% mannitol. (**D** to **H**) Box plot summary of (D) primary root total area, and cell-type specific areas of (E) epidermis, (F) cortex, (G) endodermis, and (H) pericycle cells from transversal cross sections of the early differentiation zone in WT and *kor1-4* 5-do seedlings, grown in the absence or presence of 3% mannitol. Medians are represented inside the boxes by solid lines, circles depict individual cell measurements from 10 roots. Letters denote statistically significant differences among samples as determined by ANOVA followed by Tukey’s HSD test (P < 0.05).

### Heightened JA levels guide *kor1* roots towards greater water availability

The activation of JA signalling often leads to an induction in defense responses accompanied by a reduction in organ growth (*37*). It is thus possible that increased JA production in *kor1* contributes to slow down the mutant’s root growth rate to better adjust to cellulose deficiency, and concomitantly constitutively protect the mutant from potential threats found in the soil. However, blocking JA production in *kor1-4* did not alter root elongation nor the expression of defense marker genes, as the mutant had the same root growth rate as its JA-deficient *kor1-4 aos* counterpart and similar low abundance of *VSP2* and *PDF1*.*2* transcripts (Fig. 6A, Fig. S7A and B). What could then be the consequence of increased JA signalling in *kor1*?

**Fig. 6.**
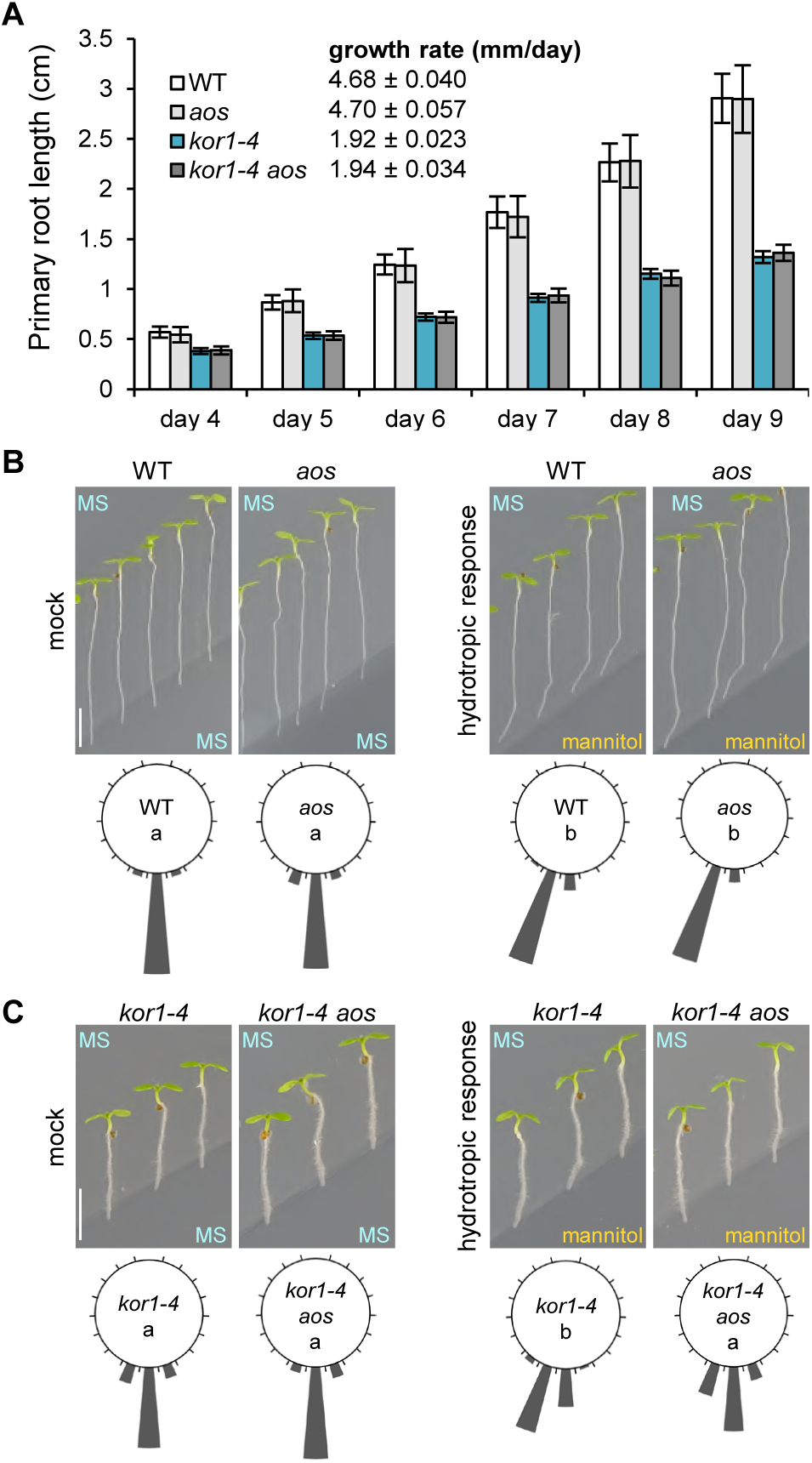
Increased JA production facilitates *kor1* response to root hydrotropism. (**A**) Primary root elongation of indicated genotypes between 4- and 9-days post germination. Bars represent the means of 40-50 plants. Data were used to determine the root growth rate in mm per day by linear regression. (**B** and **C**) Root hydrotropic response of (B) WT and *aos*, and (C) *kor1-4* and *kor1-4 aos* seedlings. Representative images and circular histograms summarizing root curvatures of indicated genotypes 24 h after transfer to split-agar Murashige and Skoog (MS) plates under mock (MS/MS) or hydrotropism-inducing (MS/400 mM mannitol) conditions. Bars indicate the percentage of seedlings exhibiting a root bending angle assigned to one of the 18 20° sectors on the circular diagram (individual measurements are available in Fig. S7D and E), n = 42. Letters indicate statistically significant differences as determined by Two-Way-ANOVA followed by Tukey’s HSD test (P < 0.05). Scale bar, 5 mm.

As the stunted *kor1-4* root growth was alleviated by continuous growth in mannitol (Fig. 5C), we tested if the mutant might preferentially grow towards hyperosmotic conditions in a split-agar assay used for measuring root hydrotropic responses among genotypes elongating at similar growth rates (*38*). Following transfer to split-agar plates with mannitol harbouring asymmetric water availability, both WT and *aos* seedling roots readjusted their growth away from the hyperosmotic media in search for greater water availability (Fig. 6B, Fig. S7C and D). Contrary to our expectations, *kor1-4* roots also bent away from mannitol with a positive hydrotropic root curvature despite their slower growth rate. To our surprize, the JA-deficient *kor1-4 aos* double mutant failed to effectively readjust its root growth direction towards a greater water availability and instead grew into the mannitol media (Fig. 6C, Fig. S7E). This effect was not a consequence of the double mutant’s inability to undergo general root bending responses such as those triggered by root gravitropism, but rather to a specific insensitivity to root hydrotropism (Fig. S7F to H). Our data thus unveiled that the constitutive activation of JA-Ile production and signalling in *kor1-4* serves to facilitate water foraging.

**Fig. S7.**
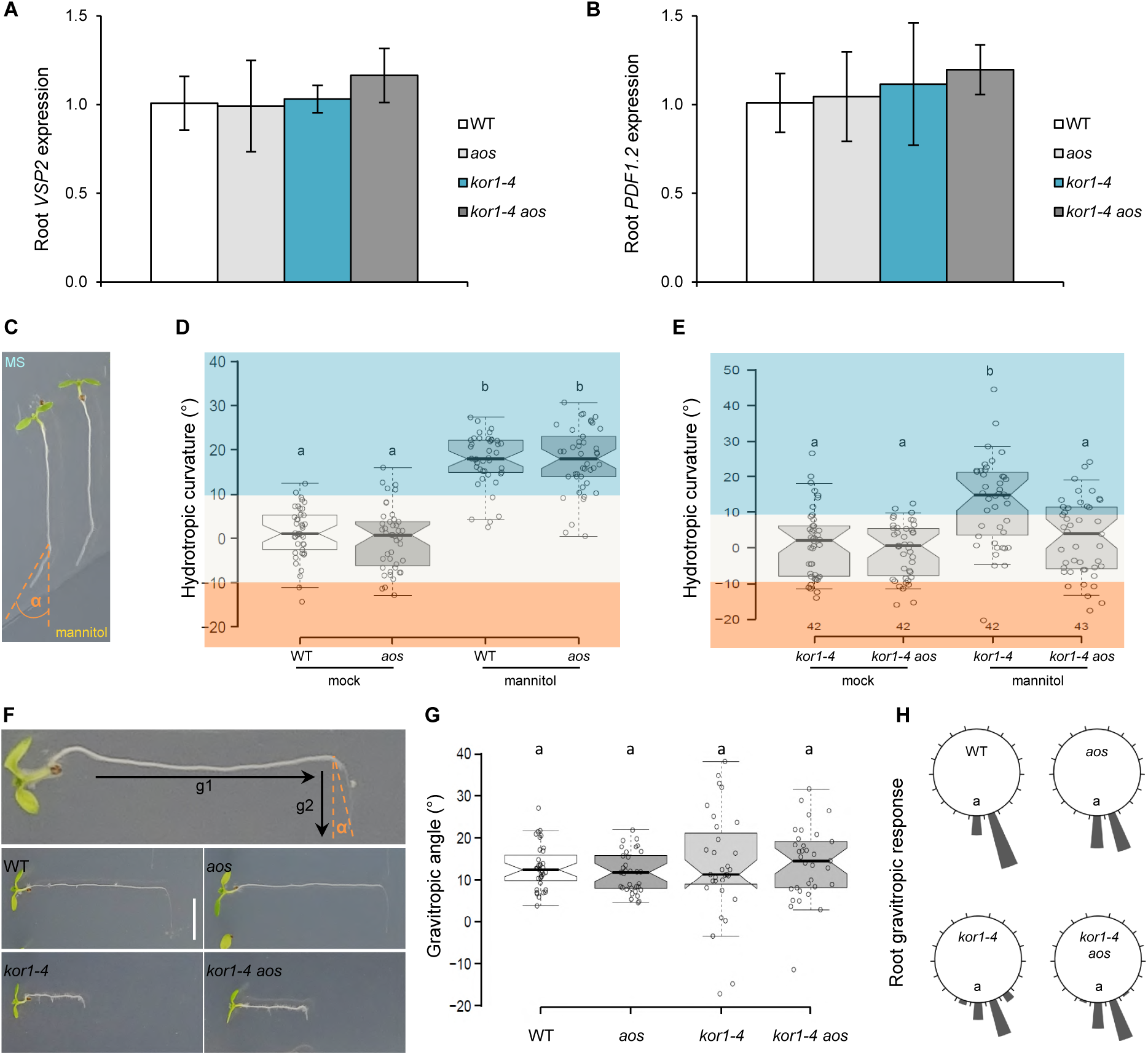
Elevated JA levels in *kor1* do not alter defense marker gene expression nor root gravitropism. (**A** and **B**) qRT-PCR of (A) *VSP2* and (B) *PDF1*.*2* basal levels in roots of indicated genotypes. Transcript levels were normalized to those of *UBC21* and displayed relative to the WT controls. Bars represent the means of 3 biological replicates (±SD), each containing a pool of ∼60 organs from 5-do seedlings. (**C** to **E**) Hydrotropic root curvature in seedlings grown with similar growth rates: WT and *aos*; and *kor1-4* and *kor1-4 aos*. (C) Representative image of a WT seedling 24 h after transfer to a split-agar plates under hydrotropism-inducing (MS/400 mM mannitol) conditions depicting the measurement of its hydrotropic root curvature angle (α). Box plot summary of root hydrotropic curvatures in (D) WT and *aos*, and (E) *kor1-4* and *kor1-4 aos* seedlings 24 h after transfer to split-agar (MS/MS, mock) or hydrotropism-inducing (MS/400 mM mannitol) plates. Medians are represented inside the boxes by solid lines, circles depict individual measurements (n = 42). Hydrotropic responses were highlighted as positive (blue, towards greater water availability), neutral (light grey, straight growth) and negative (orange, towards lower water availability) along the y-axis. (**F** to **H**) Root gravitropic response of indicated genotypes. (F) Representative images depicting seedlings grown vertically for 5-d in the 1^st^ gravity direction (g1), turned by 90° and grown for additional 24 h in the 2^nd^ gravity vector (g2) before measuring the gravitropic root curvature angle (α). (G) Box plot summary of the root gravitropic angle of WT, *aos, kor1-4* and *kor1-4 aos* seedlings. Medians are represented inside the boxes by solid lines, circles depict individual measurements (n = 31-35). (H) Circular histograms summarizing root growth curvatures of indicated genotypes from (G). Bars indicate the percentage of seedlings exhibiting a root bending angle assigned to one of the 18 20° sectors on the circular diagram. Letters in (D, E, G, and H) indicate statistically significant differences as determined by Two-Way-ANOVA followed by Tukey’s HSD test (P < 0.05).

## DISCUSSION

Plant cell walls are defining plant structures which serve a wide range of biological functions including shaping cell morphology, providing mechanical support for growth, and connecting cells within tissues. Changes in their structure and assembly may have pronounced repercussions on intracellular as well as whole-plant responses (*21*). Here we investigated how cues derived from perturbed cell walls integrate with the production of the stress hormone JA-Ile. The identification of cellulose-deficient *kor1* alleles displaying elevated JA-Ile levels specifically in seedling roots prompted us to examine cell wall-triggered induction of hormone biosynthesis in cell-type-specific contexts. By restoring KOR1 function in specific layers of the primary root, we showed that ectopic JA-Ile signalling in endodermal and pericycle cells was caused by cell non-autonomous signals deriving from adjacent and markedly enlarged cortex cells. Swollen *kor1* cortex cells could exert an increased mechanical pressure upon both external epidermal cells as well as inner tissues. As JA-Ile production is readily triggered by mechanical stress, why was the hormone increase observed only in inner tissues? A possibility is that epidermal cells could dissipate the cortex-exerted mechanical pressure by expanding towards the available outer space and by stimulating cell division. Mechanical cues are known to influence cell cycle progression in animals, and although this has not been demonstrated in plants yet, they guide the orientation of plant cell division planes (*20, 39*). Contrary to outer tissues, endodermal and pericycle cells are physically constrained and may become mechanically compressed by enlarged cortex cells. Indeed, reducing the swollen *kor1* cortex cell size in three independent manners fully abolished increased JA-Ile signalling in inner tissues. First, restoring KOR1 activity in the cortex complemented both cortex cell size and JA-Ile marker gene expression. Second, we identified a genetic suppressor of *kor1* in which cortex cell size and JA levels were also recovered. Third, by reducing *kor1* turgor pressure with hyperosmotic treatments, enlarged cortex cells and JA signalling were fully recovered. Collectively, the data demonstrate that the JA pathway is triggered by turgor pressure-driven mechanical changes in response to cellulose-deficiency. Our findings might explain why wounding single-cells by laser ablation in living roots does not induce JA-Ile signalling (*27*). The disintegration of a target cell provokes adjacent cells to bulge towards the vacuity left by the ablated cell, in a process compatible to turgor pressure changes in adjacent cells due to the vanished support from the damaged cell (*40, 41*). However, similarly to epidermal cells in *kor1*, wound-adjacent cells are not subjected to compression and might be able to compensate the increased mechanical stress by expanding towards the void space without activating JA production. On the other hand, mechanical wounding and chewing insect herbivores squash tissues and possibly compress spatially constrained cells adjacent to wounds. Because cells within tissues are connected through their cell walls, mechanical stress signals arising from turgor pressure changes may propagate over distances depending on the extent of damage (*42*).

Consistently, it has been proposed that mechanical tissue damage causes sudden pressure changes in vascular xylem vessels, compressing cells adjacent to vessels where JA biosynthesis is then initiated (*24, 43*). Hence, mechanical compression generating turgor pressure changes may be a crucial elicitor of JA-Ile biosynthesis in circumstances extending beyond cell wall perturbations, with putative osmo- and mechano-sensors located across different subcellular compartments (*44*). However, measuring mechanical stress patterns in vivo, particularly from cells embedded in inner tissues which are not physically accessible, remains challenging (*45*). Emerging tools are offering encouraging prospects to assess mechanobiological processes in vivo in future studies (*46*).

In our search to identify genetic regulators of ectopic JA-Ile production in *kor1* with potential roles in stress signals arising from turgor pressure changes, we did not retrieve known components involved in osmoregulation such as mechanically activated ion channels OSCA1 or MSL10, nor cell wall integrity sensors such as THESEUS1 whose loss of function was insensitive to isoxaben-induced JA production (*25, 47-49*). This might be due to the screen not being saturated or to genetic redundancy. Instead, the *kor1* suppressor screen recovered an allele in *ESMD1*, a putative *O*-fucosyl-transferase involved in pectin homeostasis (*31*). Although ESMD1’s substrate and biochemical function are unknown, it was proposed that it may *O*-fucosylate cell wall integrity sensor proteins and thereby contribute to cell signalling (*31*). Recently, two enzymes belonging to the same GT106 family were shown to be rhamnosyltransferases involved in the biosynthesis of the pectin component rhamnogalacturonan I (RG-I) (*32, 50*). Given that our monosaccharide composition analysis in genotypes with *esmd1-3* showed a substantial rhamnose reduction which in Arabidopsis derives mostly from RG-I (*51*), and in agreement with other GT106 family members, it is possible that ESMD1 is directly involved in pectin biosynthesis. Although it remains unclear how *esmd1* restores cell shape and JA-Ile levels in *kor1*, RG-I can have significant and specific contacts with cellulose microfibrils (*52*). As turgor pressure is counterbalanced by cell walls being under tension (*53*), it is possible that the diminished cell wall tensile strength in *kor1* is compensated by lack of *ESMD1*. The identification of ESMD1 also indicates that cell size and mechanical compression on inner tissues can be alleviated by remodelling cell wall properties and thus re-equilibrate turgor pressure.

A remarkable observation was that constitutive JA-Ile signalling enabled *kor1* roots to modify their growth towards greater water availability. This finding highlights the importance of characterizing cell wall mutants as they may provide opportunities to uncover more subtle functions of the JA pathway. While the root hydrotropic response of the JA-deficient *aos* mutant did not differ from the WT, JA-deficiency in *kor1* prevented roots bending away from the media with reduced water availability. These findings imply that while JA-deficiency *per se* does not impact root hydrotropism, the constitutive activation of JA-Ile signalling in precise tissues and cell types is beneficial for root water foraging. Importantly, the constitutive activation of JA signalling in *kor1* did not result in typical growth-inhibition responses (*54, 55*), suggesting that manipulating JA biosynthesis in specific cell-types might confer acclimation advantages without hampering overall growth. Interestingly, hyperosmotic mannitol treatments are also used to study plant responses to water deficiency, and applying exogenous JA to various plants renders them more tolerant to drought, reviewed in (*56*). Accordingly, Arabidopsis mutants impaired in JA production or signalling, are more drought-susceptible (*57*). Although it is unknown how constitutive JA signalling guides *kor1* roots away from mannitol, future studies deciphering the molecular mechanisms governing this effect could have beneficial impacts on breeding programs aimed at increasing plant drought tolerance.

## MATERIALS AND METHODS

### Plant material and growth conditions

*Arabidopsis thaliana* ecotype Columbia (Col) was used for all experiments. *JAZ10p:GUS* (*JGP*) transgenic lines in Col (WT) and in the JA-deficient *aos* mutant background were described previously (*6*). *JAZ10p:NLS3xVEN* was as in (*27*) but cloned and crossed into *aos* de novo. *kor1-6* (SALK_075812) was obtained from NASC and crossed into *JGP* and *aos JGP. esmd1-1* was described in (*31*). For in vitro assays, seeds were sterilized and stratified 2 days at 4 °C in the dark. Seedlings were grown on 0.5x solid Murashige and Skoog (MS, Duchefa) medium supplemented with 0.5 g/L MES hydrate (Sigma) and 0.7% or 0.85% plant agar (AppliChem) for horizontal or vertical growth, respectively. Horizontally grown seedlings were germinated on a nylon mesh placed on top of the MS media as described (*6*). Controlled growth conditions were set at 21°C under 100 µE m^−2^ s^−1^ light, with a 14 h light/10 h dark photoperiod. For soil propagation, transformation and crossing, plants were grown with the same temperature (T) and light intensity, but under continuous light.

### Histochemical detection of GUS activity and lignin deposition

GUS stainings were performed as described (*58*) and seedlings were photographed with a Leica M165 FC stereomicroscope fitted with a Leica MC170 HD camera. Lignin deposition was visualized by submerging seedlings in acidified phloroglucinol solution (1% phloroglucinol in 18% HCl) for 5 min, washing in 1x PBS buffer, mounting in 10% glycerol, and imaging using DIC optics on a Leica DM6B microscope fitted with a Leica DMC6200 camera.

### Plant treatments

Single cotyledon wounding of seedlings and MeJA (Sigma) treatments to assess *JGP* reporter activity were performed as described (*6*). To evaluate the JA response of the *JAZ10p:NLS-VEN* reporter in individual seedlings grown vertically, primary roots were mounted in mock or 10 µM MeJA 0.5x MS with 30 μg/ml propidium iodide (PI, Sigma) solution, imaged immediately (t=0) and after 2 h (n = 10). For osmotic support experiments, plant growth media was supplemented with either 3% mannitol (165mM, J&K Scientific), 3% sorbitol (165mM, Carl Roth), 3% PEG6000 (Serva) or 3% plant agar (AppliChem).

### Gene expression analyses

RNA extraction and qRT-PCR of *JAZ10* (At5g13220), *JOX3* (At3g55970), *VSP2* (At5g24770) and *UBC21* (At5g25760) was performed as described (*29, 58*). Similarly, the expression of *JAZ3* (At3g17860) was assayed with GACTCGGAGCCACAAAAGC and TACGCTCGTGACCCTTTCTTTG, and of *PDF1*.*2* (At5g44420) with TTTGCTGCTTTCGACGCAC and GCATGATCCATGTTTGGCTCC primers. RT-PCR of *KOR1* was performed with the primer pair AGATGCTGAAGCCAGAGCAG and TGTCATGGAGAGGTAATTCTGG.

### JA-Ile quantification

5-do roots were excised beneath the collet region and flash frozen to yield approximately 50 mg FW for each biological replicate. Extraction and quantitative measurements were performed as described (*29*). The limit of JA-Ile quantification (LOQ = 3x limit of detection) was determined from an Arabidopsis matrix as 0.49 pmol/g FW.

### Cloning and generation of transgenic lines

All transcriptional and translational reporter constructs were generated by double or triple Multisite Gateway Technology (Thermo Fisher). ENTRY plasmids containing cell-type specific promoters (pEN-L4-*IRT1p*-L3, pEN-L4-*PEPp*-L3, pEN-L4-*SCRp*-L3, and pEN-L4-*WOLp*-L3) were described in (*30*) and obtained from NASC. pEN-L4-*JAZ10p*-R1, pEN-L1-*NLS-3xVEN*-L2 and pEN-R2-*CIT*-L3 were as in (*6, 58*). CIT and mTurquoise (mT) fluorophores were subcloned into pDONR221 or pDONR-P2R-P3 to obtain pEN-L1-*CIT*-L2 and pEN-R2-*mT*-L3. Promoters were amplified from WT genomic DNA with oligonucleotides containing adequate restriction sites for *KOR1p* (GATGATGCTCTCTGATAAAGC and AAGTCTTTTGGGAGCTGCAA, 2.132 kb) and *ESMD1p* (ATCGACAGATCTCAATCTC and GGACGAGGACATCCTTGGTA, 2.168 kb) and cloned into pUC57 to create pEN-L4-*promoter*-R1 clones, as described (*58*). Coding DNA sequences of *KOR1* (ATGTACGGAAGAGATCCATG and TCAAGGTTTCCATGGTGCTG) and *ESMD1* (ATGCTAGCGAAGAATCGG and GGTGGCAGGAGGTGGTCTC) were amplified from WT cDNA with oligonucleotides specified in parenthesis containing appropriate *att* sites and recombined with pDONR221 or pDONR-P2R-P3 to obtain pEN-L1-*KOR1*-L2, pEN-L1-*ESMD1*-L2, and pEN-R2-*KOR1*-L3. For transcriptional reporters, pEN-L4-*JAZ10p*-R1 and pEN-L4-*ESMD1*-R1 were recombined with pEN-L1-*NLS-3xVEN*-L2 into pEDO097, as described (*58*). pEN-L4-*KOR1p*-R1 was also recombined with pEN-L1-*KOR1*-L2 into pEDO097 for complementation analysis. For translational reporters, pEN-L4-*KOR1p*-R1 or cell-type specific promoters were recombined with pEN-L1-*CIT*-L2 and pEN-R2-*KOR1*-L3 into a modified pH7m34gw vector named pFR7m34gw, which harbours seed RFP expression (*OLE1p:RFP*) instead of Hg resistance for in planta selection, to generate *promoter:CIT-KOR1* constructs. Similarly, to obtain *ESMD1p:ESMD1-mT* and *ESMD1p:ESMD1-CIT*, we recombined pEN-L4-*ESMD1p*-R1, pEN-L1-*ESMD1*-L2 and pEN-R2-*mT*-L3 or pEN-R2-*CIT*-L3 into pFR7m34gw (in both cases, fluorescence signals were undetectable, although the constructs were functionally complementing the mutant phenotype). All constructs were verified by Sanger sequencing, and transgenic plants were generated by floral dip with *Agrobacterium tumefaciens* strain GV3101. Transformed seeds expressing RFP in T_1_, T_2_ and T_3_ generations were selected by fluorescence microscopy, and segregation analysis was performed in >12 independent T_2_ lines. A minimum of two independent T_3_ transgenic lines were used for each construct to perform experiments and verify reproducibility.

### Confocal microscopy

Confocal laser scanning microscopy was performed on Zeiss LSM 700 or LSM 880 instruments. For live imaging, 5-do vertically-grown seedling roots were mounted in 0.5x MS with 30 μg/ml PI. As *kor1* roots are thick and recalcitrant to PI penetration, *kor1* genotypes were fixed in 4% paraformaldehyde, cleared with ClearSee, and stained with Direct Red 23 (Sigma) as described (*59*). Excitation/ detection ranges were set as follows: VENUS (VEN) and CITRINE (CIT): 514/ 515-545 nm; mTurquoise (mT): 458/ 460-510 nm; Direct Red 23: 514/ 580-615 nm; Propidium iodide (PI): 561/ 600-700 nm. All images shown within one experiment were taken with identical settings, and by analyzing at least 10 individuals per line. Image processing was performed in Fiji. Z-Stacks were displayed as texture-based volume renderings using the 3DViewer plugin of Fiji.

### Suppressor screen and mapping by NGS sequencing

Approximately 5 000 seeds (0.1 g) of *kor1-4 JGP* were mutagenized with ethyl methanesulfonate (EMS, Sigma) as described (*6*). Resulting M_1_ plants were either harvested individually (n = 1 243) or in pools of 12 (n = 230). 20 M_2_ seedlings were screened from individually harvested plants, and 480 M_2_s were screened from each pool to enlarge both the screening breadth and depth. A total of 135 260 M_2_ seedlings, from 4 003 M_1_ plants, were assayed for lack of *JGP* activity in 5-do *kor1-4* seedlings by live GUS staining as described (*6*). To increase the screen stringency and avoid the recovery of false positives, M_2_ seedlings were shifted from 21°C to 26°C 24 h prior GUS staining as *kor1* mutants are known exacerbate their phenotypes at higher T (*15*). Putative M_2_ suppressors were transferred to soil and crossed to JA-deficient (*aos, opr3, jar1*) and JA-insensitive (*coi1*) mutants to avoid the recovery of expected genes, and back crossed to *kor1-4 JGP* for segregation analysis, phenotype confirmation, and mapping population development. *esmd1-3* was identified as a *JGP* suppressor of *kor1-4* by pooling 120 individuals lacking *JGP* reporter activity from an BC_1_F_2_ population and sequencing the bulk segregants by whole genome sequencing. Genomic DNA extraction was as in (*6*). Library preparation (Illumina Shotgun TruSeq DNA PCR-free) and Illumina sequencing on a HiSeq X platform with a 150bp PE read length was performed by Macrogen. The output of 15Gb resulted in an average of 118 sequencing depth for each base across the genome. EMS-generated SNPs were identified as described (*6*), with updated software tools. Sequence reads were mapped to the TAIR10 *Arabidopsis thaliana* genome with bowtie2 aligner (v. 2.3.1, parameter-end-to-end). Alignment files were converted to BAM with SAMtools (v 1.8), and SNPs calling was performed with GATK tool (v.4.1.0.0). Common SNPs with the *kor1-4 JGP* parental line were filtered out with the intersectBed tool from BEDTools utilities (v.2.22.1). The SNPEff tool (v.2.0.4 RC1) was used to predict the effect of the SNPs in coding regions. SNP frequencies (the number of reads supporting a given SNP over the total number of reads covering the SNP location) were extracted using the Unix command awk and plotted with R. Candidate SNPs were identified in genomic regions with high SNP frequencies (0.5-1) linked to the causal SNP, which had the expected frequency of 1. Validation of the candidate SNP was done by allelism test and by complementation by transformation.

### Cell wall composition analysis

Alcohol-insoluble residue (AIR) was extracted from shoots and roots of 12-do seedlings as previously described (*60*). Each biological replicate consisted of ∼100 shoots (∼150 mg FW) or ∼300 roots (∼80 mg FW). A slurry solution of AIR (1 mg/mL water) was prepared for each sample and homogenized using a ball mill followed by sonication. Matrix polysaccharide composition of 300 µg of AIR after 2 M trifluoroacetic acid hydrolysis was analyzed via high-performance anion-exchange chromatography with pulsed amperometric detection (HPAEC-PAD), similar to (*60*), but on a 940 Professional IC Vario ONE/ChS/PP/LPG instrument (Metrohm) equipped with Metrosep Carb2 250/4.0 analytical and guard columns. Each run consisted of neutral sugar separation (22 min; 2 mM sodium hydroxide and 3.4 mM sodium acetate isocratic gradient), followed by uronic acid separation (23 min; 100 mM sodium hydroxide and 170 mM sodium acetate), and re-equilibration (14 min; starting eluents) steps.

Cellulose was quantified based on the two-step sulfuric acid hydrolysis method described by (*61*), with some modifications. Aliquots of AIR (200 µg each) were first pretreated with concentrated sulfuric acid (to swell cellulose) or were directly used for Seaman hydrolysis (to measure non-crystalline glucose), using ribose as an internal standard. Hydrolysed glucose was quantified using the HPAEC-PAD system described above but with a shorter run: 2 mM sodium hydroxide and 3.4 mM sodium acetate isocratic gradient (22 min), followed by a 3 min rinse with 80 mM sodium hydroxide and 136 mM sodium acetate, and a 4 min re-equilibration with starting eluent.

### Sectioning, segmentation and cell analysis

Roots from vertically-grown 5-do seedlings were vacuum infiltrated and fixed in glutaraldehyde: formaldehyde: 50mM sodium phosphate buffer (pH7.2) (2:5:43, v/v/v) for 1 h, dehydrated through an EtOH series, and embedded in Technovit 7100 resin (Heraeus Kulzer) as described (*58*). Samples were sectioned on a Microm HM355S microtome with a carbide knife (Histoserve) into 5 μm sections, mounted in 10% glycerol, and cell walls were visualized under dark-field of a Zeiss AxioImager microscope fitted with an AxioCam MRm camera. TIF images were segmented with PlantSeg (*62*) using preset parameters of the prediction model “lightsheet_unet_bce_dice_ds1x”, which empirically segmented our images most accurately. For display purposes only, dark field images were inverted and segmented images re-coloured in Photoshop to visualize different cell types more easily.

### Root growth and tropism assays

Primary root length was evaluated in 7-do seedlings as described (*6*) and root growth rate was determined by measuring primary root length in 4-do seedlings for 6 consecutive days. Root diameter was assessed in 7-do seedlings by imaging vertically grown seedlings on a Leica M165 FC stereomicroscope fitted with a Leica MC170 HD camera, measuring the root thickness in the early differentiation zone (marked by the appearance of root hairs). Root gravitropism assays were performed on vertically-grown 5-do seedlings by rotating the plates by 90° and evaluating root bending angles 24 h after the rotation on scanned images with Fiji. For root hydrotropism assays, 5-do seedlings were transferred to split-agar plates containing either mock (MS/ MS) or 400 mM mannitol (MS/ mannitol) by aligning root tips 3 mm from the split-media boundary (*63*). Root bending angles were evaluated 24 h after transfer to split-agar plates, by analyzing scanned images with Fiji as described (*63*).

### Statistical analysis

Box plots, multiple comparisons [analysis of variance (ANOVA) followed by Tukey’s honest significant difference (HSD) test], and circular histograms were performed in R.

## ACKNOWLEDGEMENTS

We thank Edward E. Farmer for enabling the initiation of this work by gifting the *kor1-4* and *kor1-5* EMS alleles originally identified in his lab, and for critical comments on the manuscript. We also thank Grégory Mouille for sharing *esmd1-1* seeds, Ivan F. Acosta for gifting the pFR7m34gw plasmid, Christine Wagner for assistance with AIR sample preparation, Bo Yang for HPAEC-PAD maintenance, Claus Wasternack and members of the Gasperini lab for critical discussions.

## FUNDING

This work was supported by the Leibniz Institute of Plant Biochemistry from the Leibniz Association, the German Research Foundation (DFG grant GA2419/2-1) to D.G. and (DFG grant 414353267) to C.V., and a Marie Skłodowska-Curie postdoctoral fellowship to M.K.M.

## AUTHOR CONTRIBUTIONS

S.M. and D.G. designed research; S.M, M.Z., M.K.M. and D.G. performed research; S.M., M.K.M., R.D., and D.G. analyzed data; H.S. and B.H. quantified JA-Ile levels; C.V. performed cell wall composition analyses; S.M. and D.G. wrote the manuscript with input from all authors.

## COMPETING INTERESTS

The authors declare no competing interests.

## Notes

### Competing Interest Statement

The authors have declared no competing interest.

